# Resolving Local and Global Conformational Heterogeneity of the Human Intrinsically Disordered Proteome

**DOI:** 10.64898/2025.12.16.694679

**Authors:** Hossain Shadman, Jesse Dylan Ziebarth, Qianyi Cheng, Mohamed Laradji, Yongmei Wang

## Abstract

Linking the sequences of intrinsically disordered regions (IDRs) to their structural ensembles and biological functions remains a central challenge in understanding how disorder encodes activity. A recent study has shown that human IDRs with different levels of compactness, as measured by Flory’s exponent, are associated with specific cellular functions and localizations, and Flory’s exponent can be predicted from sequence features with reasonable accuracy. However, IDRs are known to sample highly heterogenous conformations that can be masked by ensemble-averaged metrics such as Flory’s exponent. Here, we introduce a simple framework that resolves heterogeneous conformations. We pair two polymer physics descriptors, shape ratio (*R*_s_) and relative shape anisotropy (RSA), to construct joint two-dimensional (RSA, *R*_s_) maps at both the global and local (subchain) scales. We show that sequences with similar Flory’s exponent can display strikingly different two-dimensional conformational maps that reflect differences in charge patterning. IDRs with similar global maps can exhibit markedly different local maps that can be linked to local sequence variations. This leads to the identification of a class of IDRs that appear non-compact globally but contain locally compact subchains. These locally compact IDRs are found to be associated with similar GO functional and cellular localization enrichments and phase-separation propensities as globally compact IDRs. Our framework moves beyond ensemble-averaged descriptors, providing new tools that capture the intrinsic heterogeneity of IDR conformations and thus offer new opportunities to link IDR sequences with functions.

**Significance Statement:** Intrinsically disordered proteins (IDPs) and regions (IDRs) exhibit highly heterogeneous conformational ensembles, yet studies routinely use ensemble-averaged metrics to characterize them that obscure this heterogeneity. We present a simple, general framework that quantifies structural heterogeneity in human IDRs. This resolves forms of structural heterogeneity that are otherwise overlooked and enhances our understanding of how IDR structures relate to their sequences and functions. A python package for immediate implementation of this framework is provided.

## Introduction

The three-dimensional structures of proteins exist on an order-disorder continuum, ranging from stable, folded regions, such as α-helices, to disordered regions with diverse conformational landscapes known as intrinsically disordered regions (IDRs) and proteins (IDPs).^1,2^ These disordered regions are present in ∼70% of human proteins.^3^ IDPs/IDRs mediate cellular signaling processes, drive liquid-liquid phase separation, and have been linked to significant human conditions, such as aging, Alzheimer’s disease, mental conditions (e.g. ADHD) and cancer.^3–13^ However, the sequence-structure-function relationships of these crucial members of the human proteome are only beginning to be understood.^13–18^ To help fill this gap, Tesei *et al.*^19^ recently performed molecular simulations of 28,058 IDRs from ∼15,000 distinct full-length proteins in the human proteome that were selected using pLDDT confidence scores from AlphaFold2.^20,21^ One of the key results of this work was the establishment of links between the conformational properties (e.g., the Flory scaling exponent, *ν*) of human IDRs and their functions. Both compact and extended IDRs were associated with specific cell functions and localizations, with compact IDRs, for example, associated with chromatin binding and signaling receptor activity, and localization to nuclear bodies, endosomes, and mitochondria. In addition to insights into sequence-structure-function relationships, this work provides a rich data source for further analysis of the human intrinsically disordered proteome.

The conformational properties of IDPs/IDRs have been commonly characterized using ensemble averages.^3,15,22^ However, the conformations of IDRs are usually highly diverse and dynamic, and ensemble averaging can mask this intrinsic structural heterogeneity. Here, we re-analyze the human disordered proteome dataset (Tesei *et al.*^19^) – examining both instantaneous values and ensemble averages – to help resolve the heterogeneity of human IDRs. Since IDRs are polymer chains, we use polymer descriptors to demonstrate that ensemble averages can obscure substantial conformational heterogeneity. For example, IDRs that would be classified as extended, through ensemble average quantities, adopt compact conformations. Additionally, two IDRs can achieve very similar ensemble-averaged quantities (*ν*) through different modes of interplay in their underlying polymer properties. By analyzing polymer properties at both the global (i.e. whole chain/region) and local^15^ (subchain) scales, we demonstrate that IDRs can display locally compact regions that are not apparent from global descriptors. Notably, proteins containing locally compact but globally non-compact IDRs exhibit Gene Ontology (GO) functional and cellular localization enrichments, and intrinsic phase-separation propensities similar to proteins with globally compact IDRs. Our work reveals subtle and previously overlooked aspects of IDP/IDR conformations and, thus, offer insights that can aid future investigations into their sequence-structure-function relationships, and enhance future protein design^23–25^ strategies.

## Materials and Methods

### I: Data Source for the Human Intrinsically Disordered Region-ome

Tesei *et al.* used AlphaFold2 pLDDT confidence scores to select the human IDR-ome: a set of ∼28000 Intrinsically Disordered Regions (IDRs) from the human proteome.^19^ They conducted coarse-grained simulations using the CALVADOS 2 package^26^ of each of these ∼28000 human IDR sequences, generating trajectories of 1000 snapshots per simulation. The IDR-ome trajectories, along with the associated conformational properties and sequence features are publicly available on their website (https://sid.erda.dk/cgi-sid/ls.py?share_id=AVZAJvJnCO) and GitHub repository (https://github.com/KULL-Centre/_2023_Tesei_IDRome).^19^ We downloaded their trajectories and sequence/structure data for analysis in June 2024 and used MDTraj^27^ to analyze trajectories. Conformation visualization was done using VMD.^28^ Four IDRs (‘Q53SF7_218_1128’, ‘Q7Z2Y5_341_1224’, ‘Q9Y2W1_1_611’, ‘Q9BXT5_1_968’) were excluded from our analysis due to their inconsistent data sizes, resulting in a dataset of 28,054 IDRs.

### II: Simulations of Model Polymers (GW, SAW, PEI)

We simulated a Gaussian walk (GW) polymer model chain using a previously described^29^ method to generate ∼1,000,000 snapshots of a 100-monomer chain. Briefly, the GW chain was simulated in continuum space (i.e. not on a lattice), where the displacement of each monomer from the previous one, along the x-, y-, and z-directions, was drawn from a Gaussian distribution with a mean (μ) of 0 and a standard deviation (σ) of 1 unit. No additional constraints were imposed on the model. In this study, the GW provides a reference landscape for the conformational landscapes of other peptides/polymers.

We also conducted Monte Carlo simulations of a self-avoiding walk (SAW) polymer model chain consisting of 100 monomers on a simple cubic lattice, as described previously.^29^ Excluded volume interactions were enforced such that no two monomers could occupy the same lattice site. We conducted 10 independent runs using different random seeds, with each run generating ∼35,000 SAW conformations. These 10 runs were combined to generate ∼350,000 total snapshots. In each run, the initial ∼10% of the configurations, representing the transient period before equilibration, were discarded.

Our group previously conducted all-atom molecular dynamics (MD) simulations of a 40-mer polyethyleneimine (PEI) chain at different protonation states (0% protonation or 0p to 100% protonation or 40p).^30^ Briefly, the PEI chain at each protonation state was simulated separately in a box with periodic boundary conditions, containing TIP3P water and salt ions. For each protonation state, one trajectory comprising 45,000 snapshots (spanning 90 ns) was generated. Here, we re-analyze the results of these PEI simulations.

### III. Polymer Physics Quantities: R_s_, R_g_, RSA

We compute the instantaneous shape ratio^29,31^ (*R_s_*), defined as:

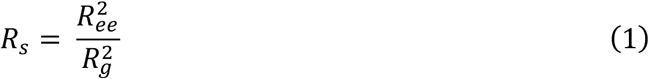

where *R_ee_* and *R_g_* are the end-to-end distance and radius of gyration, respectively. Lower *R_s_* values (∼2 or lower) characterize collapsed chains, whereas fully extended chains have *R_s_* ∼12. Additionally a chain can adopt *R_s_* > 12 by minimizing *R_g_* and maximizing *R_ee_*, as we demonstrated previously.^29^ In this study, 〈*R*_*s*_〉 is computed as 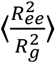 and not as 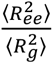.We also compute the relative shape anisotropy (RSA) using gyration tensors as described previously.^29^ RSA can range from 0 (perfectly isotropic structure) to 1 (straight, thin rod).^29,32^ Snapshots of GW conformations at different (RSA*, R_s_*) combinations are shown in Figure S1 (Supplemental Information). Both *R_s_* and RSA are unitless quantities.

In this study, we generate scatter plots and heatmaps^33^ of *R_s_* versus RSA, an approach that extends one we described previously,^29^ where we characterized the conformational ensembles of disordered chains by generating scatter plots of *R_s_* against fluctuations in size (computed as *R_g_*/〈*R*_g_〉). The heatmaps are based on a two-dimensional histogram of the *R_s_* against RSA scatter plots (using numpy.histogram2d, with density set to ‘True’, which computes the probability density function as bin count / sample count / bin area) with 200 bins along the RSA*-*axis and 100 bins along the *R_s_*-axis. We implement a Gaussian filter (scipy.ndimage.gaussian_filter with sigma = 1) on the two-dimensional histograms to generate the heatmaps.

Throughout this study, *ν* refers to the Flory scaling exponent. For the IDR sequences, we use *ν* values previously computed and published by Tesei *et al.*^19^ However, we do compute *ν* for IDR subchains (covered later), using the same *ν* formula as Tesei *et al.* (as provided through their GitHub account). Briefly, an IDR’s *ν* value was computed by Tesei *et al.* as follows: for all residue pairs (*i*, *j*) separated by more than 5 residues along the sequence (|*i* − *j*| > 5), *ν* was obtained by fitting the relationship 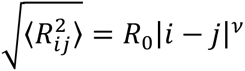 to the root-mean-square residue-residue distances 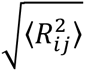.

To quantify the relative sizes of polymers, we compute an expansion factor 〈*⍺*〉 which is defined by:

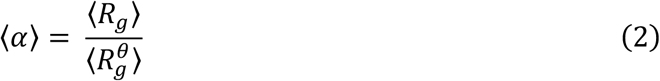

where 〈*R*_g_〉 is the mean radius of gyration of a polymer and 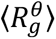 is its mean radius of gyration in a θ-solvent, when the monomer–monomer and monomer–solvent interactions are balanced. For a polymer in a θ-solvent, 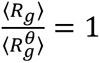 and *ν* = 0.5. For 〈*⍺*〉 > 1, the monomer-solvent interaction is stronger than the monomer-monomer interaction and the chain is expanded (i.e., *ν* > 0.5). In contrast, 〈*⍺*〉 < 1 signifies a poor solvent that not only can lead to chain collapse but also liquid-liquid phase separation. For the IDRs considered in the present work, we approximate 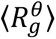 using the Analytical Flory Random Coil (AFRC^34^) python package (https://afrc.readthedocs.io/en/latest/). AFRC computes a distribution of *R_g_*^θ^ values for any peptide chain, based solely on the amino acid sequence. For each human IDR, we compute a single 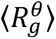 from its AFRC-generated *R_g_*^θ^ distribution. We then calculate 〈*⍺*〉 using Eq. (2) for that specific IDR. We note that 〈*⍺*〉 has been referred to in previous^15^ work as normalized *R_g_* or 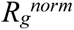.

### IV. Sequence Features

The sequences of amino acid residues in IDRs enable the calculation of various sequence features unique to each IDR. For each IDR sequence, we compute the net charge per residue (NCPR) as follows:

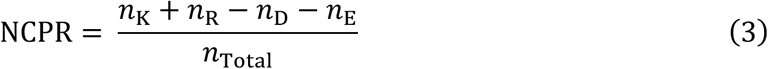

where *n*_K_, *n*_R_, *n*_D_and *n*_E_ represent the numbers of lysine, arginine, aspartic acid and glutamic acid residues, respectively, and *n*_Total_ is the total number of amino acid residues in the sequence. Other IDR sequence features that were calculated by Tesei *et al.* are used in parts of our analysis, such as in constructing random forest regressor models (detailed later). The sequence features we used are: SHD, SCD, NCPR, *z*(δ_+–_), *f_aro_*, 〈*λ*〉, κ, *z*(Ω_–_), *f_neg_*, *f_pos_*, FCR, *z*(Ω_+_) *z*(Ω_h_), *f_P_*, and *z*(Ω_π_). The definition and significance of these sequence features can be found in earlier studies.^35–40^ Most of these were defined and computed by Tesei *et al.*^19^ FCR is the fraction of charged residues. κ, SCD (sequence charge decoration), and *z*(δ_+–_) (z-score to estimate the segregation between positive and negatively-charged residues) are parameters that describe the charge distribution along the chain. SHD, which is the sequence hydropathy index, is a parameter that describes the patterning of hydrophobic residues. 〈*λ*〉 is a residue ‘stickiness’ parameter that measures the strength of attractive intra-chain interactions relative to solvent-protein interactions. *f_aro_* and *f_P_* represent the fractions of aromatic residues and proline, respectively.

### V. f_C_ and f_C_shape_ scores

For each IDR simulation, we compute the *f_C_shape_* score, which is an adaptation of the *f_C_* score that we defined previously.^29^ The *f_C_* score quantifies the conformational diversity of a given polymer/protein by comparing its structural heterogeneity to that of a reference GW chain. The *f_C_* score of a given polymer/protein was calculated by analyzing a scatter plot of *R_s_* versus *R_g_*/〈*R*_g_〉 comparing that polymer/protein to a reference GW chain. Likewise, *f_C_shape_* is defined from the comparison of a scatter plot of *R_s_* versus RSA of a polymer with that of the reference GW chain. The *f_C_shape_* score, like the *f_C_* score, ranges from 0 to 1 and quantifies the conformational diversity of a given polymer/protein by measuring the fraction of the reference GW conformation space visited by that polymer/protein. Specifically, we first transform the data (details provided previously^29^) presented on an (RSA, *R_s_*) scatter plot, and compute the *f_C_shape_* score as:

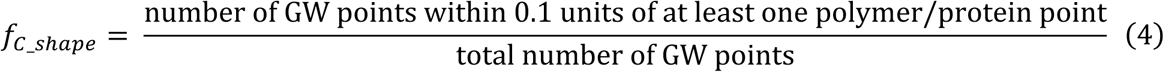

Like *f_C_* scores in our previous^29^ study, disordered proteins are expected to have *f_C_shape_* scores at least an order of magnitude higher than structured proteins, though different *f_C_shape_* scores are expected for different disordered proteins. Figure S2 shows that *f_C_* and *f_C_shape_* are highly correlated with each other in the human IDR-ome dataset (Pearson correlation coefficient = 0.706).

### VI. IDR Subchain Analysis

We examine IDR conformations at the local level using subchain^15,33^ analysis. We use a moving/sliding window across each IDR chain and calculate quantities such as *R_s_* and RSA for those subchains. We note that, for an IDR simulation trajectory, quantities (e.g. 〈RSA〉) computed over the entire chain are global quantities, whereas those computed over a portion (or ‘subchain’) of an IDR are local quantities. For instance, a moving window of size 3 over an example chain “*ABCDEFGH*” considers the subchains *ABC*, *BCD*, *CDE* etc.

In our analysis, we use a moving window of size 30 for all IDRs with more than 60 residues. For IDRs with fewer than 60 residues, our moving window sizes are 1/3^rd^ of the number of residues in the respective IDR chain. This moving window size was chosen because the smallest IDR in the human IDR-ome dataset is 30 residues long. To evaluate the local conformational properties of a given IDR, we compute quantities such as 〈RSA_*i*_〉 and 〈*R*_*s*,*i*_〉 for each moving window *i* over that IDR’s sequence. Based on those local conformational properties, we compute two new quantities for each IDR: SM_RSA and SM_*R_s_*. For each IDR, SM_RSA is defined as:

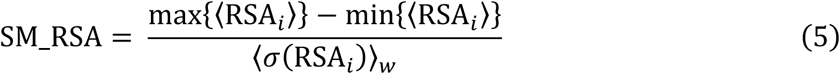

Where max{〈RSA_*i*_〉} is the maximum value of 〈RSA_*i*_〉, and min{〈RSA_*i*_〉} is the minimum value of 〈RSA_*i*_〉, with *i* varying between 1 and the total number of moving windows of that IDR. For each moving window *i*, the standard deviation of its RSA, *σ*(RSA_*i*_), is computed. 〈*σ*(RSA_*i*_)〉_<_ is the average of all values of *σ*(RSA_*i*_) across all windows of the IDR. SM_*R_s_* is computed similarly as shown below:

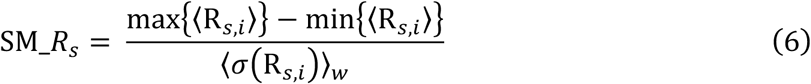

SM_RSA and SM_*R_s_* are positive and capture how local conformations fluctuate for a given IDR, or ‘local smoothness.’ Lower SM values (<1), suggest few local conformational fluctuations, whereas higher SM values indicate higher local conformational fluctuations.

### VII. Machine Learning Models

For all ∼28000 IDRs in the human IDR-ome dataset, we implement random forest regressor models and use sequence features to predict 〈RSA〉, 〈*R*_*s*_〉, 〈*⍺*〉, *ν*, SM_RSA and SM_ *R_s_*. We use the scikit-learn^41^ library in python (using the RandomForestRegressor command) to implement our models, on 50:50 train-test splits of the dataset. We examine a scatter plot of actual vs predicted values for a single test set. We also combine metrics of the test sets from 10 different randomly generated train-test splits. For these 10 test sets, we generate combined feature importance plots, reported as mean ± standard deviation.

### VIII. Gene Ontology (GO) Enrichment Analysis of IDRs with Varying Compactions

Molecular Function (MF) and Cellular Component (CC) Gene Ontology (GO)-term lists for the proteins corresponding to the human IDRs were pre-processed and published by Tesei *et al.*^19^ We downloaded two files from their GitHub repository (https://github.com/KULL-Centre/_2023_Tesei_IDRome): *df_GO_molfunc.pkl* and *df_GO_cellcom.pkl*, containing MF terms for ∼22,000 IDRs and CC terms for ∼25,000 IDRs respectively. We used these two files to assess functional and cellular localization enrichment of IDR subgroups with varying compactions, treating MF and CC as separate categories of GO terms. Using an IDR’s overall (global) *ν* value, its *ν_min_local_* (minimum local *ν*), and *ν_max_local_* (maximum local *ν*), we defined four subgroups (Table S1). Subgroups 1 and 2 consist of IDRs that are globally compact (*ν* < 0.45) and globally extended (*ν* > 0.55), respectively, as suggested originally by Tesei *et al*.^19^ Subgroup 3 consists of IDRs that are globally non-compact but contain locally compact regions (*ν* ≥ 0.45 but *ν_min_local_* < 0.45). Subgroup 4 consists of IDRs that are globally non-extended but contain locally extended regions (*ν* ≤ 0.55 but *ν_max_local_* > 0.55). We note that non-compact does not necessarily mean extended (and vice versa).

We use Fisher’s exact tests (using fisher_exact from scipy.stats) to assess GO-term enrichment in each subgroup relative to its background. The MF dataset (∼22,000 IDRs) served as the background for the MF subgroups and the CC dataset (∼25,000 IDRs) served as the background for the CC subgroups. For each GO term, we constructed a 2×2 contingency table based on the number of IDRs annotated with or without that term in the subgroup and corresponding background group. We computed p-values using a one-sided test for enrichment and corrected for multiple testing with the Benjamini–Hochberg false discovery rate (FDR-adjusted p < 0.05 was considered significant) using fdrcorrection from statsmodels.stats.multitest. As some background groups had sample sizes much larger than those of their subgroups, we implemented a random subsampling approach on the background group, as follows. If the background group (MF or CC) contained more than five times as many IDRs as its subgroup, we drew 20 random subsamples of the background (each of size = 5 × subgroup sample size), and recalculated the GO enrichment of that subgroup relative to each background subsample. Only GO terms that remained FDR-significant in at least 15 out of these 20 iterations were retained in the final GO enrichment results for that subgroup.

Hardenberg *et al.* previously^42^ ranked and published *p_LLPS_* values of the proteins in the human proteome, where *p_LLPS_* is a metric that represents the intrinsic probability of a protein to mediate spontaneous liquid-liquid phase separation (LLPS). In this study we compute the mean ± standard deviation *p_LLPS_* of the proteins corresponding to the IDRs in the four subgroups (Table S2).

We note that *p_LLPS_* values and GO terms were assigned to proteins (with UniProt identifiers) and not individual IDRs. GO-term enrichment was performed at the IDR level. A caveat of this approach is that some proteins contribute multiple IDRs that inherit identical GO annotations, so individual IDR entries are not strictly independent. However, in subgroups 1 and 3, at least 94% of the IDRs map to unique proteins (Tables S1 and S2). Additionally, at most ∼11.6% of unique proteins are shared between subgroups 1 and 3 (Tables S1 and S2). Thus, any non-independence affects only a small fraction of the data. Collapsing to one IDR per protein would discard proteins contributing multiple IDRs. We report effect sizes (odds ratios, fold enrichment) together with FDR-adjusted p-values.

## Results

### I: (RSA, R_s_) is a Universal Map for Disordered Chains

We quantify the conformational ensembles of IDRs using two dimensionless descriptors of shape: instantaneous shape ratio^29,31^ (*R_s_*), the ratio of the structure’s *R_ee_*^2^ (end-to-end distance) to *R_g_*^2^ (radius of gyration), and relative shape anisotropy^29,32^ (RSA). These two metrics have a relatively low correlation (the Pearson correlation coefficient between *R_s_* and RSA is 0.269 for the conformations of the GW chain and 0.351 for the instantaneous conformations of the IDR-ome; data not shown), indicating that they capture different features of structure. When combined, (RSA, *R_s_*) provides a better description of shape than either quantity does alone (Figure S1 shows selected conformations of GW chains with varying (RSA, *R_s_*) combinations). For example, conformations with both *R*_*s*_ ≈ 12 and RSA ≈ 1 resemble fully straight rods, as expected (Figure S1c); however, if only one of these conditions is satisfied, the corresponding structures are not necessarily rod-like (Figures S1b and S1d).

Figure 1a presents (RSA, *R_s_*) scatter plots for the Gaussian walk (GW) and self-avoiding walk (SAW) polymer models. The GW plot serves as a reference landscape in these plots, as the conformations of unrestricted GW chains provide a background for those of other polymers and proteins whose conformations are limited by excluded volume and other types of interactions.^29^ The points for the SAW in the (RSA, *R_s_*) scatter plot lie within the bounds established by the GW chain (Figure 1a). The difference between SAW and GW scatter plots is relatively small; SAW covers nearly the entirety of the reference (GW) plot and has a high *f_C_shape_* score (close to 1). Figures 1b and 1c present heatmaps of the GW and SAW (RSA, *R_s_*) scatter plots, respectively. The SAW chains tend to adopt more extended conformations (i.e., the heatmap is shifted more to the upper right), consistent with the effect of excluded volume interactions in the SAW chain model.

**Figure 1:**
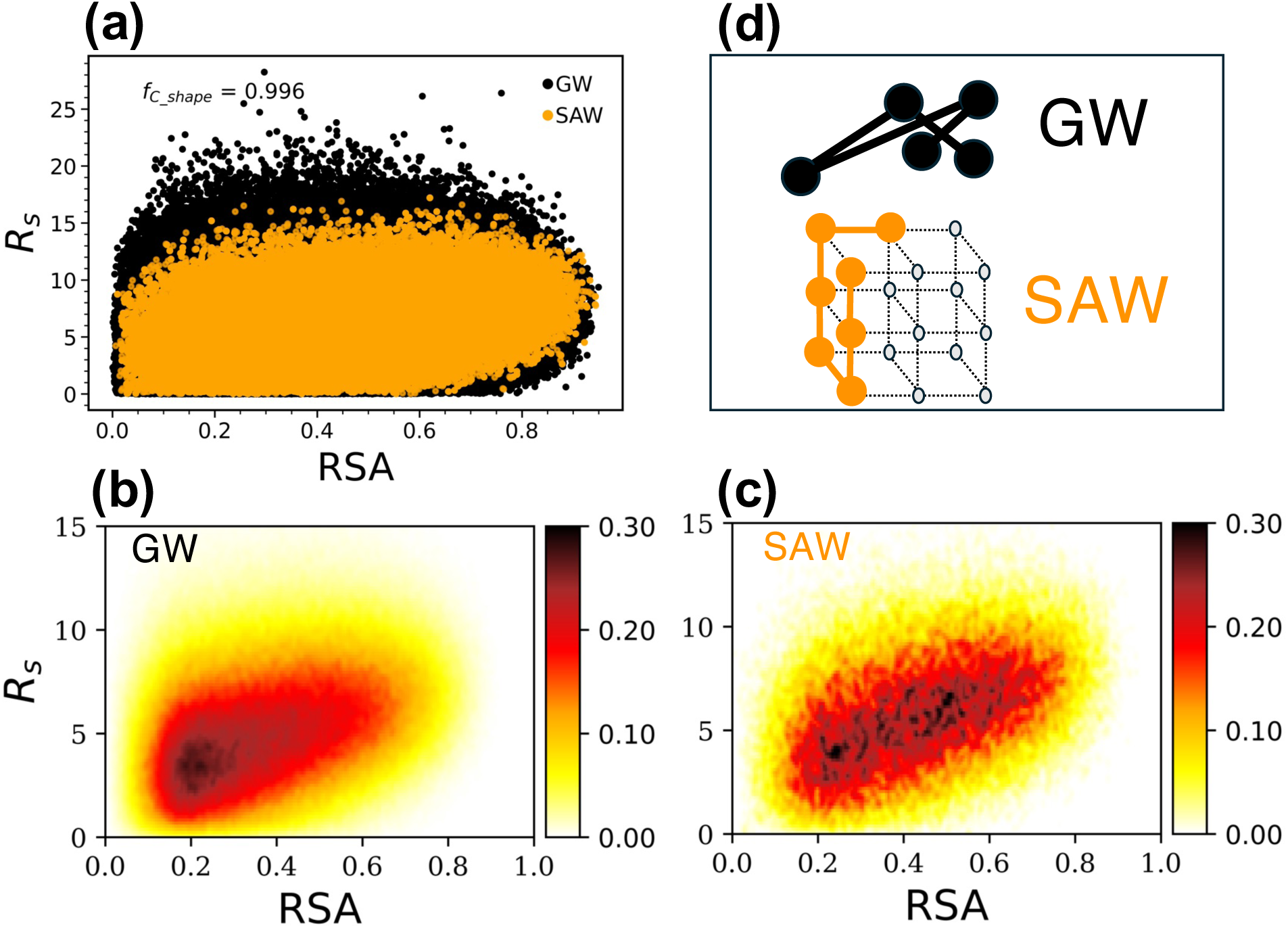
*R_s_* versus RSA of the GW and SAW polymer models. (a) GW and SAW (RSA, *R_s_*) scatter plots, (b) GW (RSA, *R_s_*) heatmap, (c) SAW (RSA, *R_s_*) heatmap. (d) Illustrative examples of GW and SAW models.

Our group previously^30^ conducted MD simulations of the nonviral gene delivery vector polyethyleneimine (PEI) at protonation states ranging from 0% protonation (0p) to 100% protonation (40p). There are dramatic differences in the conformational ensembles of PEI at different protonation states, with lower protonation states taking on collapsed conformations and higher protonation states adopting rod-like conformations.^29,30^ To visualize these differences, we examine (RSA, *R_s_*) scatter plots (Figure 2a) and heatmaps (Figure 2b). In contrast to the SAW model, PEI at most protonation states visits only a limited amount of the conformational space visited by GW. While PEI chains with relatively low protonation have relatively high conformational diversity, they are less likely to adopt conformations with high RSA and high *R_s_* values than GW chains. On the other hand, highly protonated PEI (35p and 40p) has strong intrachain electrostatic repulsion and primarily adopts extended conformations (i.e. those with high *R_s_* and RSA values). The (RSA, *R_s_*) heatmaps (Figure 2b) also clearly show the transition from collapsed to extended conformations as protonation increases, as the peaks in the heatmaps shift from lower to higher values of *R_s_* and RSA with increasing protonation. This trend is also reflected in Figure S3, where PEI has low 〈RSA〉 (∼0.2) at 0p (i.e. visiting collapsed conformations) and relatively high 〈RSA〉 (∼0.7) at 40p (i.e. preferring rod-like conformations), behavior mirroring trends in mean *R_s_* for each protonation state that we reported previously.^29^

**Figure 2:**
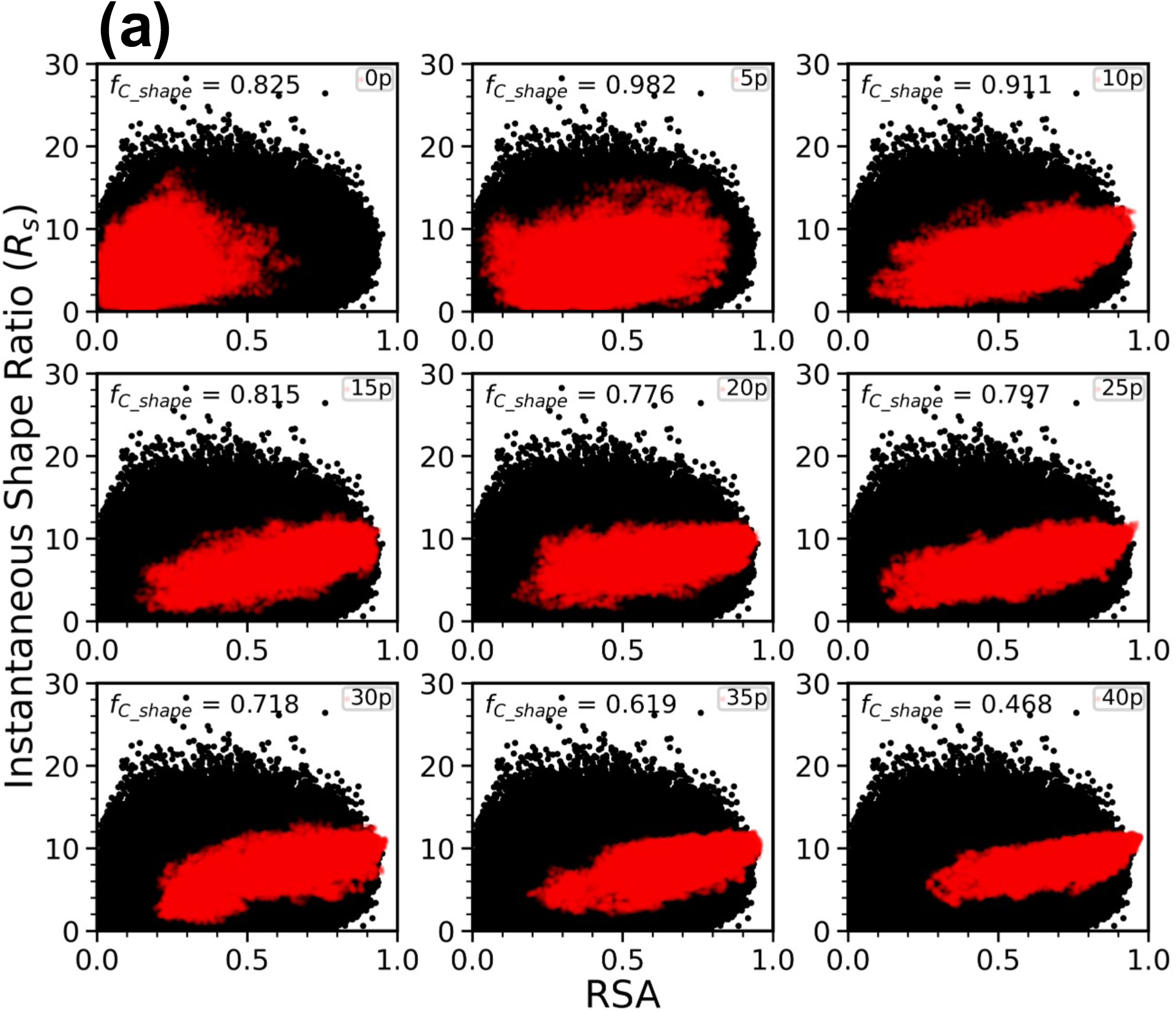
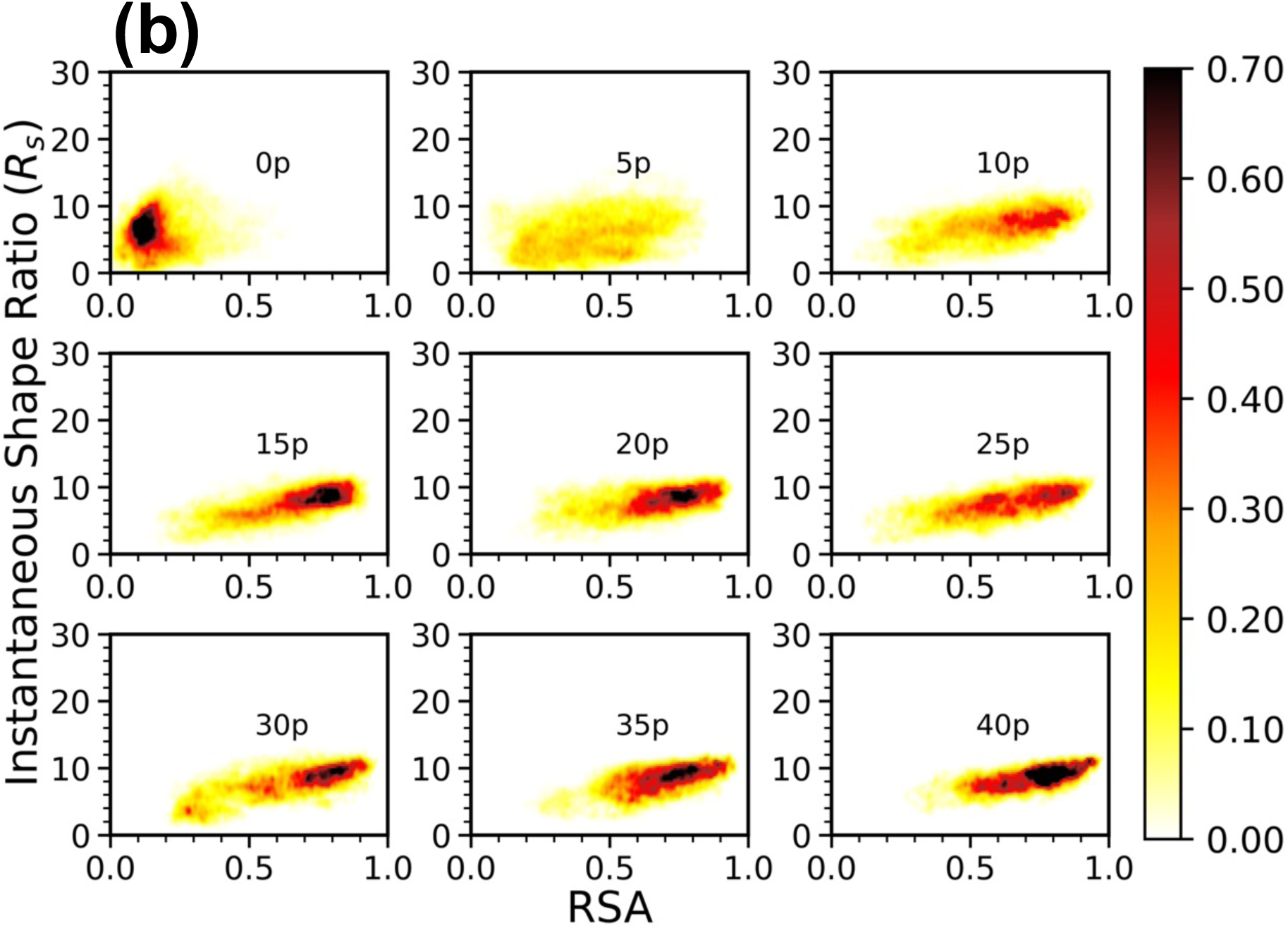
*R_s_* against RSA for a 40-mer polyethyleneimine (PEI) chain, (a) (RSA, *R_s_*) scatter plots for PEI at different protonation states along with GW for comparison (black dots: GW, red dots: PEI) (b) (RSA, *R_s_*) heatmaps for PEI at different protonation states. Protonation states range from 0p or 0% to 40p or 100%.

### II: Resolving the Heterogeneity of the Global Conformational Ensembles of the Human IDR-ome

To better understand the conformational ensembles of the human intrinsically disordered proteome, we re-analyze the human intrinsically disordered region (IDR or “IDR-ome”) dataset published by Tesei *et al.*^19^ The (RSA, *R_s_*) scatter plot of the entire IDR-ome interestingly overlaps almost completely with that of the GW reference (Figure 3a). This indicates that the diversity of the conformational ensemble created by combining the entirety of the human IDR-ome is comparable to that of the GW polymer model. This similarity is further confirmed by the closeness of their corresponding (RSA, *R_s_*) heatmaps (Figure 3b for the IDR-ome; Figure 1b for GW) and the nearly identical distributions of *R_s_* (Figure 3c) and RSA (Figure 3d) of both the IDR-ome and the GW model.

**Figure 3:**
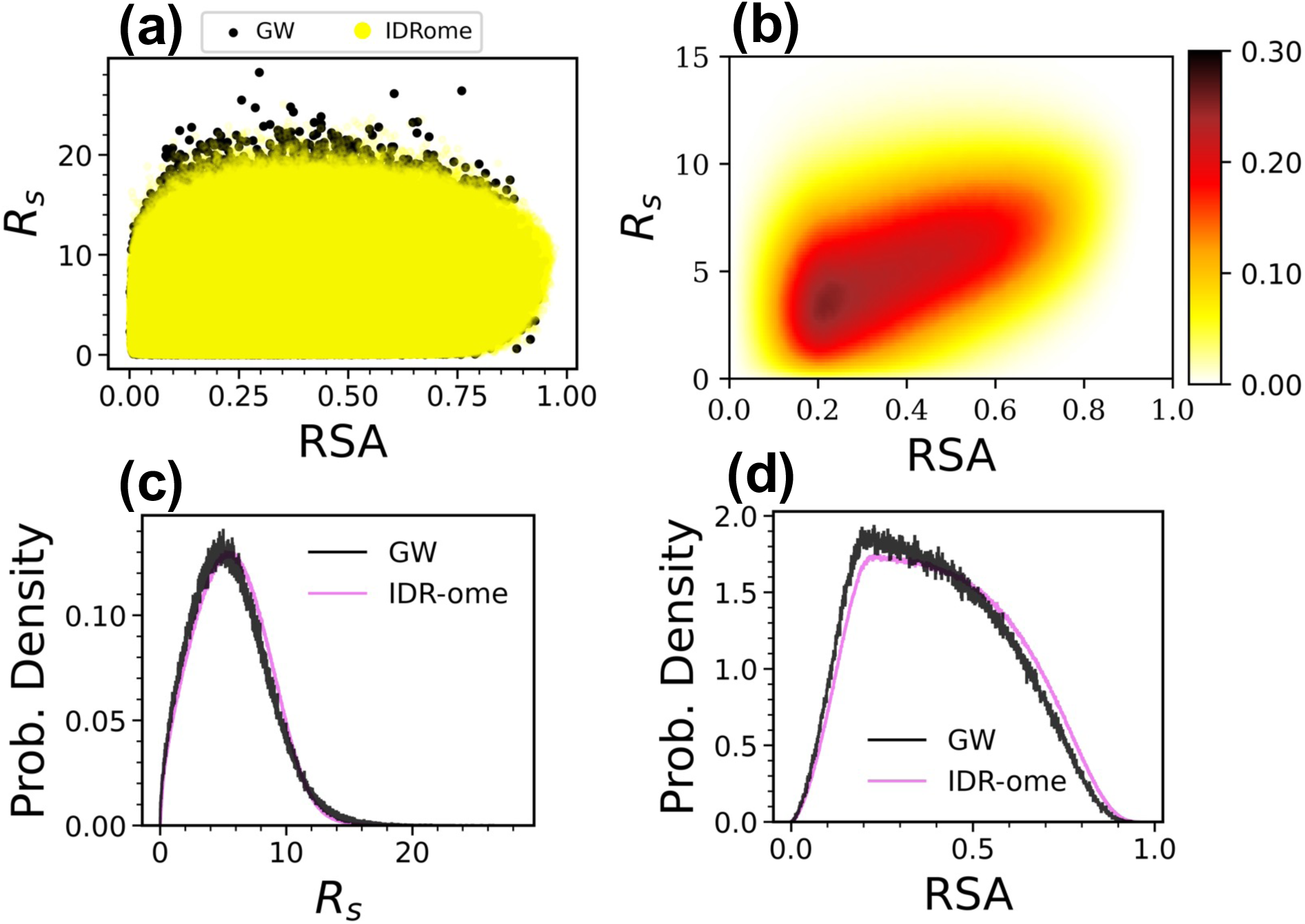
For all simulation snapshots in the human IDR-ome dataset: (a) (RSA, *R_s_*) scatter plots, (b) (RSA, *R_s_*) heatmap, (c) distribution of all *R_s_* values and (d) distribution of all RSA values. For (a), (c) and (d), data for GW is provided as a reference.

For each IDR, Tesei *et al.*^19^ computed the apparent Flory scaling exponent (*ν*), a metric that was used to define compact and extended IDRs.^19^ IDRs in the IDR-ome dataset have a wide range of *ν*, from 0.065 to 0.71, with *ν* < 0.45 indicating^19^ compact IDRs and *ν* > 0.55 indicating extended IDRs. Chain compaction, as defined by *ν*, was found to be associated with cellular function and localization.^19^ To explore how *R_s_* and RSA change with *ν*, we sorted all IDRs by *ν* and generated three (RSA, *R_s_*) heatmaps corresponding to the lowest, middle, and highest 〈*ν*〉 values, respectively, with each heatmap representing 300 sequences (Figure 4). As 〈*ν*〉 increases from compact (Figure 4a) to extended (Figure 4c) values, the highest density region of the (RSA, *R_s_*) heatmap shifts from the lower left to the upper right (compare Figures 4a, 4b and 4c). This behavior mirrors the changes in the heatmaps for PEI (Figure 2a), with higher 〈*ν*〉 in the IDR heatmaps and higher PEI protonation states both resulting in extended conformations with higher (RSA, *R_s_*) values. Importantly, all three heatmaps reveal the substantial degree of conformational diversity of the IDRs; they all show broad ranges of *R_s_* and RSA values, encompassing both collapsed and rod-like conformations, and demonstrate heterogeneity not captured by an ensemble-averaged property (*ν*).

**Figure 4:**
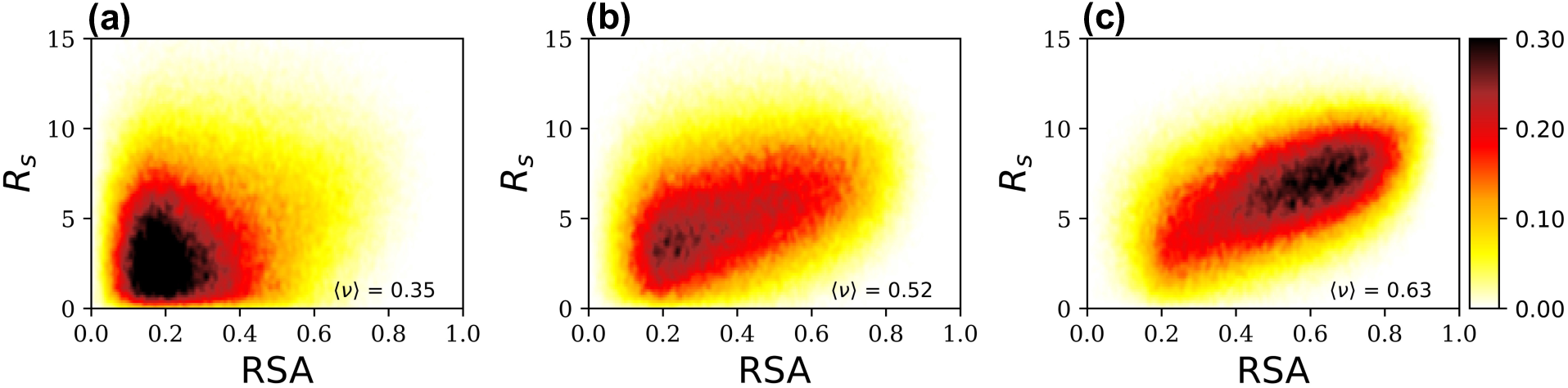
(RSA, *R_s_*) heatmaps of IDRs in the human IDR-ome with (a) low, (b) median, and (c) high 〈*ν*〉. The IDRs were first sorted by increasing *ν*; the first 300 IDRs were selected to generate the heatmap in panel (a), the middle 300 IDRs for (b) and the last 300 IDRs for (c). Each heatmap was generated from 300,000 simulation data points.

While we have, to this point, examined (RSA, *R_s_*) scatter plots created by combining many IDRs, we now turn to plots of individual IDRs. First, we compare the scatter plot of an IDR with a low *ν* value (Figure 5a; gene name: NPM2) against one with a high *ν* value (Figure 5b; gene name: SETSIP). In humans, NPM2 is involved in chromatin reprogramming, and SETSIP plays a part as a transcriptional activator.^43^ The overall trend in *R_s_* and RSA is consistent with *ν*, with points concentrated on the lower left of the plot for NPM2 (*ν* = 0.065; Figure 5a) and on the upper right of the plot for SETSIP (*ν* = 0.71; Figure 5b). Nonetheless, NPM2 also samples extended conformations (*R_s_* ∼ 11, and RSA > 0.6), while SETSIP accesses compact conformations (*R_s_* ∼ 2 and RSA < 0.2) – highlighting subtleties in each IDR’s conformational landscape not knowable from *ν* alone (Figure 5).

**Figure 5:**
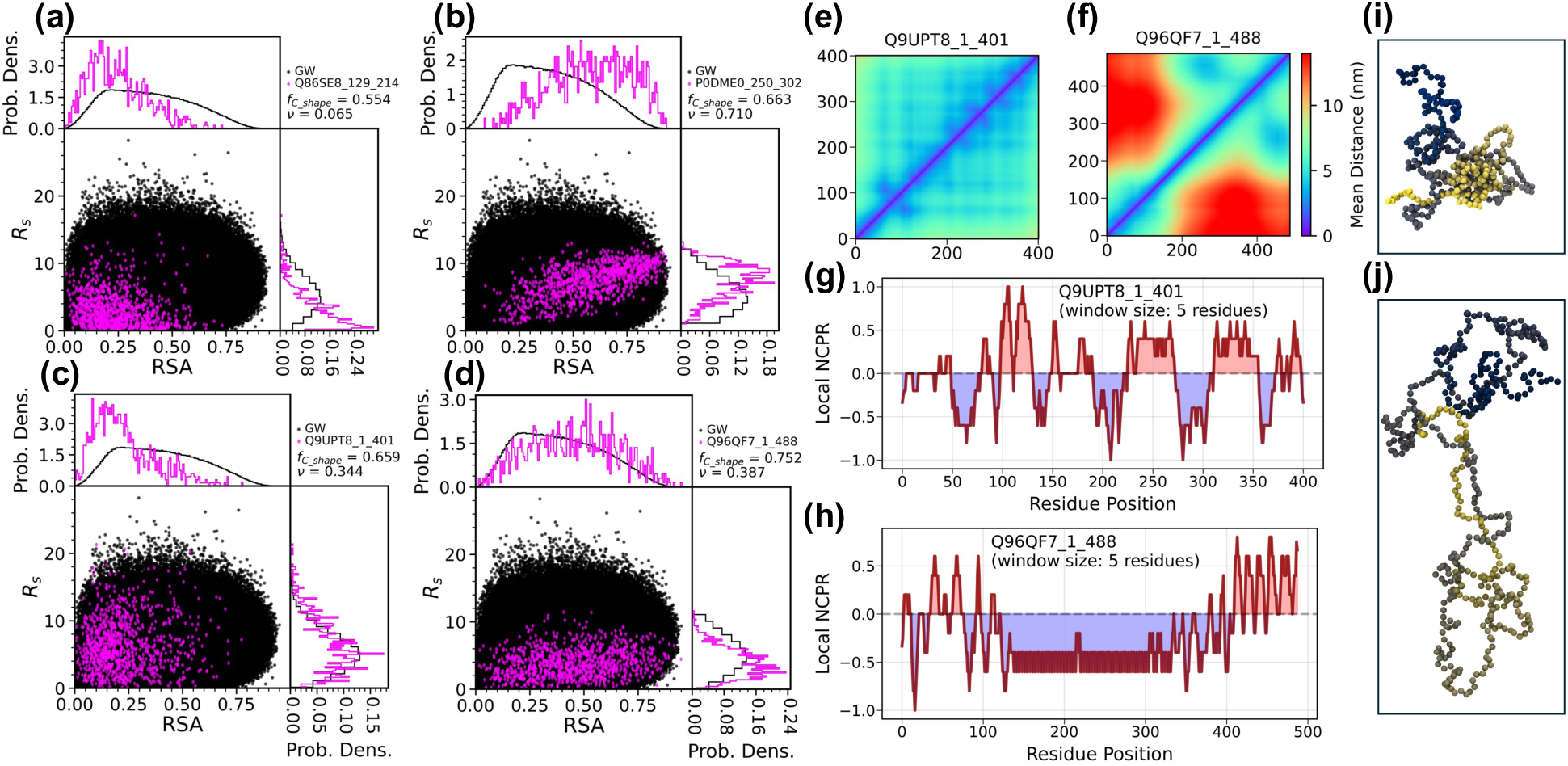
Analysis of single IDRs. Panels (a) and (b) correspond to IDRs with very low and very high values of *ν*, respectively. Panels (c) and (d) correspond to IDRs with similar values of *ν*. Panels (e) and (f) are pairwise mean distance heatmaps for the same IDRs as (c) and (d) respectively, with x and y axes showing residue indices. Panels (g) and (h) provide net charge per residue (NCPR) distributions along the sequence for the same IDRs as (c) and (d) respectively, using a moving window of 5 residues. Panels (i) and (j) show conformation images corresponding to the IDRs in panels (c) and (d), respectively. Gene names for the proteins associated with the IDRs in (a) and (b), are NPM2 and SETSIP respectively; For (c), (e) and (g), it is ZC3H4; For (d), (f), and (h), it is GCNA. The Molecular Function (MF) GO terms associated with ZC3H4 are ‘nucleic acid binding‘, ‘chromatin binding‘, ‘RNA binding‘, ‘protein-containing complex binding’, and ‘metal ion binding’ whereas for GCNA, it is just ‘other.’ The Cellular Component (CC) GO terms commonly associated with both ZC3H4 and GCNA are ‘chromosome’, ‘intracellular membrane-bounded organelle’, ‘nucleoplasm’, and ‘nucleus.’ ZC3H4 is additionally associated with CC terms ‘cytoplasm’ and ‘cytosol’ whereas GCNA is additionally associated with CC terms ‘nuclear body’ and ‘PML body’ (see Methods section for GO).

Comparison of IDRs with similar *ν* values reveals that there can be an interplay between *R_s_* and RSA distributions. Figures 5c (gene name: ZC3H4) and 5d (gene name: GCNA) show scatter plots of two IDRs with similar *ν* values (0.344 and 0.387, respectively), and both are among the most compact 2% of human IDRs (*ν* < 0.45).^19^ Additionally, both IDRs have relatively close *z*(δ_+–_) values (ZC3H4: 8.72, GCNA: 8.08), which is a metric^19^ for positive and negative charge segregation. Despite their close *ν* values, their *R_s_* and RSA distributions differ noticeably, with ZC3H4 (an RNA-binding protein^43^) accessing almost the full range of *R_s_* values but mostly low RSA (< 0.5), and GCNA accessing the full range of RSA values (indicating both spherical and rod-like conformations) but a relatively limited range of *R_s_* (< 5). Scatter plots of additional IDRs with relatively close *ν* but with subtle differences in their (RSA, *R_s_*) profiles are shown in Figure S4.

To more fully understand the differences in (RSA, *Rₛ*) profiles of the two (compact) IDRs ZC3H4 and GCNA, we examined their pairwise distance heatmaps (Figures 5e and 5f, respectively) and local net charge per residue (Figures 5g and 5h, respectively). Most residue pairs in ZC3H4 remain relatively close (< 8 nm) to one another (Figure 5e), which can be attributed to the short stretches of alternating negative and positive charge (Figure 5g), particularly in the middle of the chain, that likely promote compact sphere-like conformations (with RSA < 0.5; see Figure 5i). In contrast, a long stretch of negative charge occurs near the middle of the GCNA sequence (from residue ∼150 to ∼400; see Figure 5h), followed by a stretch of positive charge near the C-terminus. This negative charge in the middle of the chain has relatively large distance separations (>10 nm) from the residues near the N-terminus (red off-diagonal region in Figure 5f). The positive charge-dense region near the C-terminus (near residue ∼480; Figure 5h) likely interacts with this negatively charged stretch, keeping end-to-end distances relatively low, and consequently, the *R_s_* distributions (Figure 5d). This is suggested by the conformation image of GCNA (Figure 5j), which depicts a rod-like shape overall, but with the ends positioned relatively close to each other. The Molecular Function (MF) GO terms associated with ZC3H4 and GCNA do not overlap. In contrast, there are 4/6 common Cellular Component (CC) GO terms associated with both ZC3H4 and GCNA (Figure 5; see Methods for GO).

### III: Subchain-level Analysis Reveals Compact and Extended Local Structures in Human IDRs

To this point, we have characterized the conformational heterogeneity of IDRs using the properties of the entire IDR chain. However, focusing solely on these global properties ignores an important potential source of heterogeneity, as there could be significant local^15,44^ variation in the structural features along the length of an IDR. We, therefore, examined the local structures of the IDRs by using a moving window to select subchains of each IDR and analyzing the structural features of these subchains. We then computed SM quantities (SM_RSA and SM_*R_s_*) for IDR chains to measure the extent of variation in RSA and *R_s_* across subchains, with higher SM (> 1) indicating high variation in RSA or *R_s_* values across the subchains of the IDR. SM_RSA and SM_*R_s_* are largely proportional to one another in the IDR-ome dataset (Figure S5). For illustration, Figures 6a (gene name: PYDC5) and 6b (gene name: BRPF3) depict IDRs with low SM_RSA (< 0.2) and high SM_RSA (> 1), respectively. Figures S6a and S6b show SM_*R_s_* corresponding to these two IDRs. We note that PYDC5 is involved in the cellular response to viruses, and BRPF3 plays a role in the initiation of DNA replication.^43^ As illustrated in Figure 6a, IDRs with low SM_RSA have 〈RSA〉 values that are relatively stable across subchains and close to the global 〈RSA〉 value. This stability is further illustrated by the (RSA, *R_s_*) heatmaps from different subsections of PYDC5 (subA and subB; Figure 6c), which resemble one another closely, reflecting consistency in local conformational landscapes. Thus, for PYDC5, the global property is a reasonable estimate of local structural properties.

**Figure 6:**
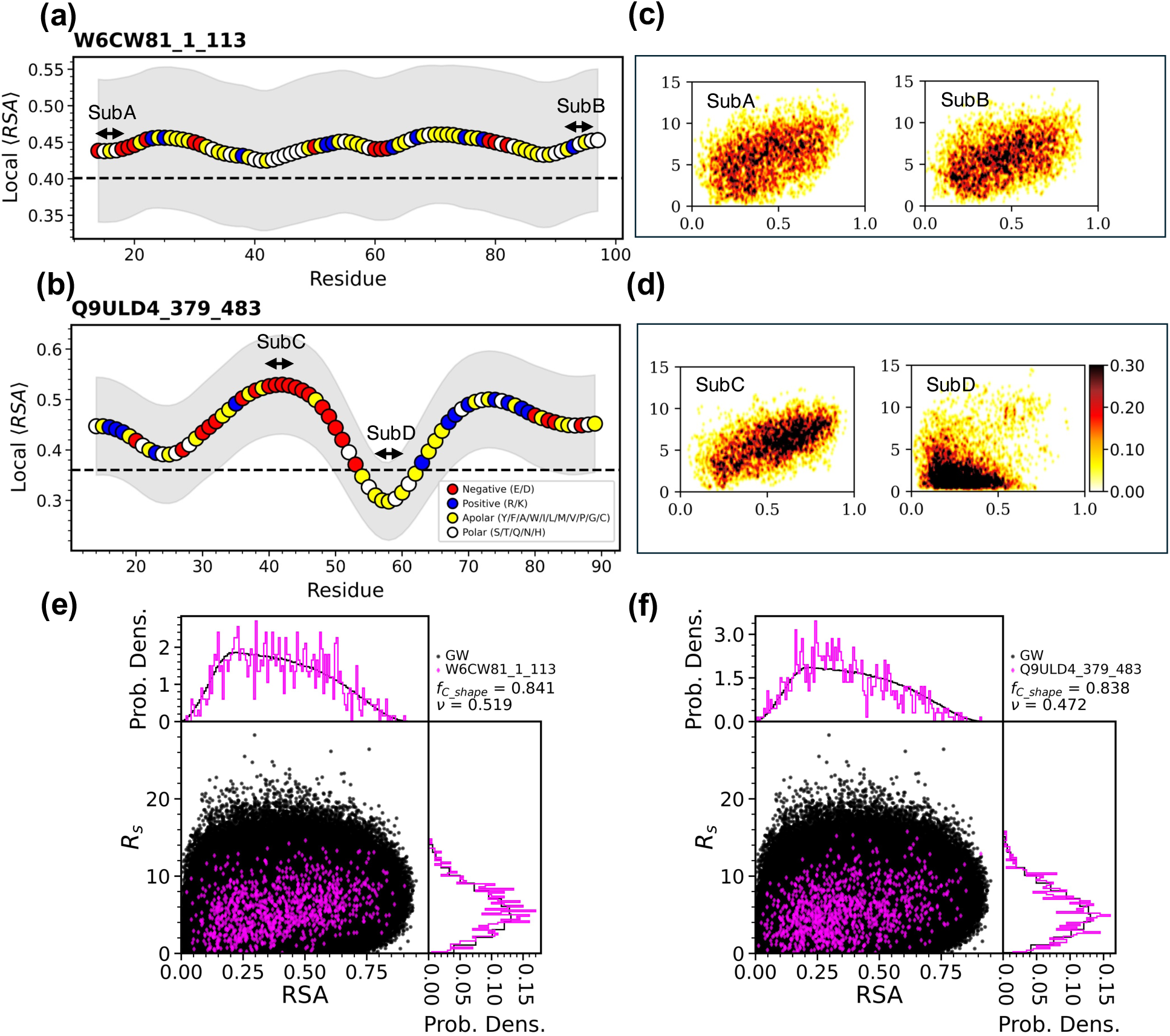
(a) and (b): Local variation in 〈RSA〉 across an IDR (identity shown in bold) chain. The horizontal axis identifies the middle residue for each moving window. The gray shaded region represents the standard deviation in RSA for the moving windows. The dashed horizontal line identifies 〈RSA〉 for the entire IDR (global). For (a), SM_RSA is 0.184 and *ν* is 0.519. For (b), SM_RSA is 1.23 and *ν* is 0.472. Panels (c) and (d) show (RSA, *R_s_*) heatmaps for select subchains, as indicated in panels (a) and (b) respectively. SubA, subB, subC, and subD have 〈*ν*〉 at 0.56, 0.56, 0.60 and 0.15, respectively. The gene names associated with the proteins for the IDRs in (a) and (b) are PYDC5 and BRPF3, respectively. (e) and (f) are global (RSA*, R_s_*) scatter plots of the IDRs corresponding to panels (a) and (b), respectively.

In contrast, there is high variability in local 〈RSA〉 in BRPF3 (Figure 6b). Some local 〈RSA〉 values of BRPF3 (e.g. subC) are almost double the global value. Additionally, as shown by the heatmaps of subC and subD (Figure 6d), subchains of BRPF3 can have conformational landscapes that are markedly different from one other (subC is quite extended whereas subD is fairly compact). Additionally, despite the dramatic differences in the local conformational landscapes of PYDC5 and BRPF3, their global (RSA, *R_s_*) scatter plots (Figures 6e, 6f, respectively) largely resemble one another, with similar *ν* and *f_C_shape_* scores. This strongly demonstrates that two IDRs with nearly indistinguishable global conformational landscapes can nonetheless exhibit very distinct local landscapes. The sequence compositions of PYDC5 and BRPF3 (Figures S6c and S6d, respectively) reveal a higher percentage of charged residues in BRPF3. The collapsed region (subD) of the BRPF3 IDR is flanked on either side by relatively long stretches of oppositely charged residues (Figure 6b). In the human IDR-ome, both SM quantities (SM_RSA and SM_ *R_s_*) can range from <1 to ∼2 or more, though most IDRs (∼97%) have SM values <1, suggesting that their local properties mirror global ones.

As global *ν* < 0.45 was used^19^ as a cutoff to define compact IDRs, PYDC5 and BRPF3 would not be considered compact, as both have global *ν* > 0.45. However, subchains of BRPF3 have *ν* < 0.45, with subD (Figure 6) having 〈*ν*〉 = 0.15. This indicates that while BRPF3 is not considered compact globally, it exhibits local compaction. Therefore, to gauge the extent of local compaction in the human IDR-ome, we computed *ν_min_local_* for all ∼28000 IDRs, where *ν_min_local_* is the minimum local *ν*. While there are only 597 IDRs with global *ν* < 0.45, there are 1742 IDRs with *ν_min_local_* < 0.45, among which 1439 IDRs have *ν* ≥ 0.45.

Globally compact IDRs were previously found^19^ to be associated with function and cellular localization. To determine whether IDRs with local compaction are associated with function and cellular localization in the same way as globally compact IDRs, we performed a GO-term enrichment analysis. Specifically, we assessed whether locally compact but globally non-compact IDRs (subgroup 3) are associated with similar GO enrichments as globally compact IDRs (subgroup 1; see Table S1 and Methods). Local compaction is relatively rare (subgroup 3 comprises ∼5% of IDRs), though it is slightly more prevalent than global compaction (subgroup 1 comprises ∼2% of IDRs), whereas local extension is more prevalent (subgroup 4 comprises ∼67% of IDRs). Using Molecular Function (MF) and Cellular Component (CC) GO-term lists previously published by Tesei *et al.*^19^ (see Methods), we compared functional enrichment across our subgroups. We first focused on the GO terms that Tesei *et al.* previously identified, using Brunner-Munzel tests, as being associated with more compact/extended IDRs, and quantified the proportion of IDRs within each subgroup annotated with those terms. In both the MF (Supplementary_Data_1.xlsx) and CC (Supplementary_Data_2.xlsx) categories, nearly all GO terms previously^19^ identified as being associated with compact IDRs (highlighted in orange) showed higher fractions of IDRs in our subgroup 1 than in subgroup 2. Moreover, for those GO terms, fractions of IDRs from subgroup 3 resembled those from subgroup 1.

We performed Fisher’s exact tests (see Methods) to determine GO-term enrichment in each CC and MF subgroup relative to its background (see Supplementary_Data_3.xlsx). MF subgroups 1 and 3 share 9/10 of their significant GO terms, and CC subgroups 1 and 3 share 8/12 of their significant GO terms, suggesting that proteins with IDRs that are globally non-compact but locally compact have similar functions and cellular localization as those containing globally compact IDRs. Though non-compact does not necessarily mean extended, subgroup 3 notably includes IDRs that are globally extended yet locally compact (*ν* > 0.55 and *ν_min_local_* < 0.45; e.g. O00566_96_359).

As chain compaction may be associated^19^ with a propensity for liquid-liquid phase separation (LLPS), we examined the mean *p_LLPS_* values (Table S2; see Methods) of the proteins containing these IDRs, where *p_LLPS_* measures the intrinsic propensity of the protein giving rise to LLPS.^42^ A threshold (*p_LLPS_* ≥ 0.61) was suggested to indicate that proteins mediate spontaneous LLPS and have droplet-driving potential under physiological conditions, though it was recognized that proteins with *p_LLPS_* values below this threshold may also mediate LLPS, if given the right conditions.^42^ IDRs in each of the four subgroups are associated with a wide range of *p_LLPS_* values (Table S2), but subgroups 1 and 3 are associated with higher mean values (〈*p*_*LLPS*_〉 ∼0.76 for both) than subgroups 2 and 4 (∼0.54 and ∼0.62, respectively). This indicates that proteins that contain globally non-compact IDRs with local compaction (subgroup 3) have the same propensity for driving LLPS as proteins that contain globally compact IDRs (subgroup 1). Notably, subgroup 3 includes an IDR from the tumor suppressor protein p53 (P04637_354_393; UniProt ID: P04637), which exists in many malignant tumors, and has been reported to undergo phase separation (*p_LLPS_* = 0.98).^42,45^ Subgroup 3 also includes two IDRs of B-Myb (P10244_1_32 and P10244_188_658; UniProt ID: P10244), which regulates cell cycle progression and cell survival and has been associated with poor patient outcomes in cancer.^46^ B-Myb also has very high *p_LLPS_* (0.98)^42^, suggesting high intrinsic potential for LLPS.

### IV: Relationships between Conformational Properties and Sequence Features

The conformational ensembles of IDRs are believed to be, at least in part, determined by their sequences, and structural properties such as 〈*R*_g_〉 and *ν* have been shown to be predictable from sequence features, such as the patterning of charged residues.^15,19,37^ Here, we use random forest models to examine how sequence features can be used to predict the ensemble-averaged properties examined in this work (〈RSA〉, 〈*R*_*s*_〉, 〈*⍺*〉, *ν*, SM_RSA and SM_ *R_s_*). With the exception of 〈*R*_*s*_〉, there is high correlation (Pearson R ≥ 0.80) between actual values and the model’s predictions (Table 1 and Figure S7), indicating that the sequence strongly influences the structure of an IDR. SCD^19^ (sequence charge decoration) and SHD^19^ (sequence hydropathy index) are the two most important sequence features in the models for 〈RSA〉, 〈*R*_*s*_〉, and *ν*, a group of structural features that are highly correlated with each other (Figure S8).

**Table 1:**
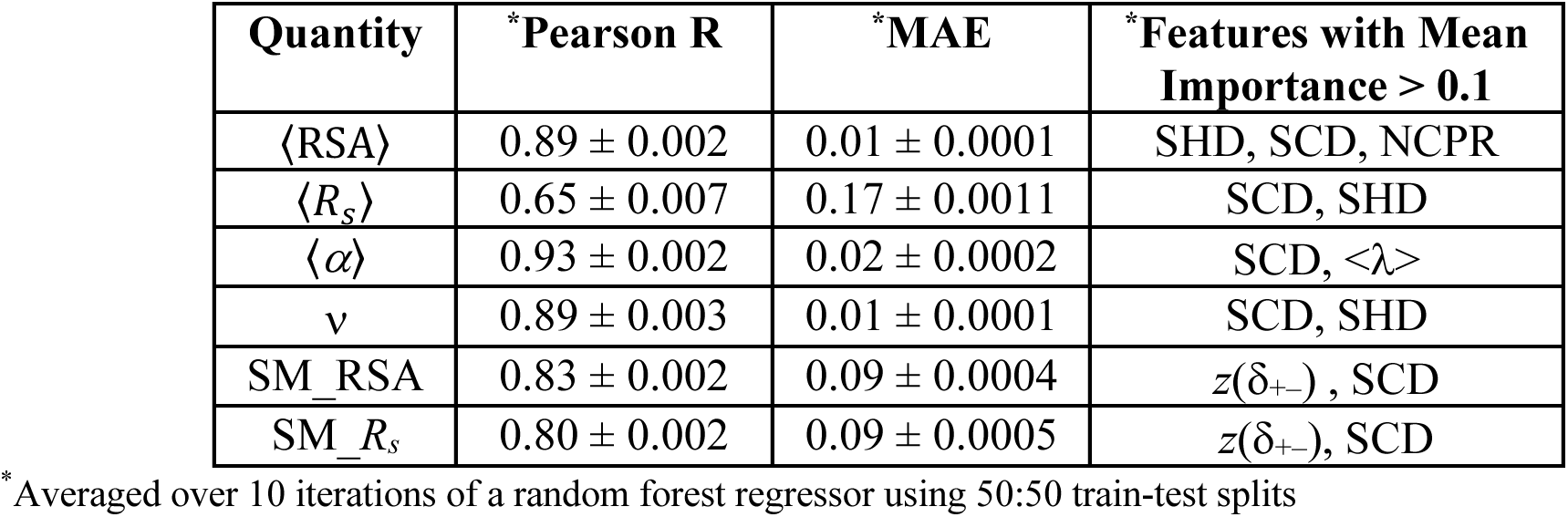
Performance and feature importance of random forest models trained on sequence features to predict ensemble-averaged descriptors for the human IDR-ome. Test-set Pearson correlations and mean absolute errors (MAE) are shown for each predicted quantity.

Both SM quantities (SM_RSA and SM_ *R_s_*) are predictable from sequence features relatively well, suggesting that subchain-level variation in conformational properties is encoded in the sequence. For both SM quantities, we observe *z*(δ_+–_) – a metric^19^ for positive and negative charge segregation – as the most important feature (Table 1 and Figure S7). SM_RSA does show some positive correlation with *z*(δ_+–_) (Figure S9a). Almost all the SM_RSA values are less than 1 for IDRs that are less than 20% charged (Figure S9b). With higher (>20%) percentages of charged residues, *z*(δ_+–_) varies widely (∼0 to 10+) and so does SM_RSA (∼0 to 1+) (Figure S9b). The IDR of BRPF3, which has high local fluctuation (SM_RSA>1; Figure 6b), is one example of an IDR with relatively high *z*(δ_+–_) (∼6). Conversely, the IDR of RBM25 is densely charged (FCR > 0.8) yet relatively stable at the subchain level (SM_RSA < 1), likely due to well-mixed positive and negative charges (*z*(δ_+–_) <1) (Figure S10).

The relationship between *ν*, 〈RSA〉, and the expansion factor 〈*⍺*〉 for the human IDR-ome is presented in Figure 7a. For IDRs with low *ν* (*ν* < 0.5), 〈RSA〉 is relatively low (mostly < 0.35). *⍺*, which represents an IDR’s *R_g_* relative to its *R_g_*^θ^ (i.e. the radius of gyration of the chain behaving as an ideal chain in a θ-solvent), is a metric commonly used in polymer physics. As a recently developed tool^34^ enables the prediction of theoretical protein/peptide *R* ^θ^ distributions directly from sequence, we computed 〈*⍺*〉 (see Methods) for all IDRs in the human IDR-ome. When *ν* < 0.5, 〈*⍺*〉 increases with *ν*, but when *ν* > 0.5, *ν* stays at a relative plateau despite continued increases in 〈*⍺*〉, resulting in IDRs with *ν* ≈ 0.5 having 〈*⍺*〉 values that range from ∼1 to ∼1.8 (Figure 7a). Most IDRs have 〈*⍺*〉 greater than 1, indicating greater 〈*R*_g_〉 relative to their corresponding 〈*R*^*θ*^〉.

**Figure 7:**
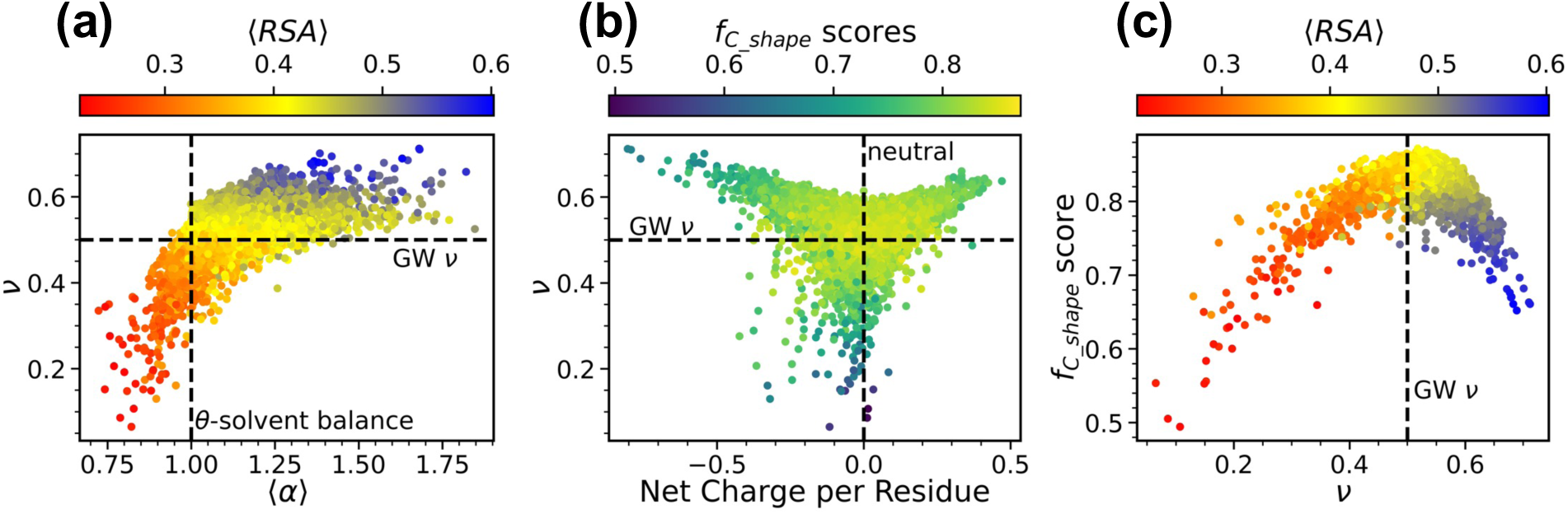
For all ∼28000 IDRs in the human IDR-ome dataset, (a) is a scatter plot of *ν* against the expansion factor 〈*⍺*〉 color-coded by 〈RSA〉, (b) is a scatter plot of *ν* against net charge per residue (NCPR), color-coded by *f_C_shape_*, and (c) is a scatter plot of *f_C_shape_* score against *ν* color-coded by 〈RSA〉. *f_C_shape_* is a measurement of the conformational diversity of an IDR.

As charged residues were previously^22,47^ shown to influence the conformational properties of disordered proteins, we explore here how net charge per residue (NCPR) is related to structural features such as *ν*, and *f_C_shape_*, a measurement of the conformational diversity of an IDR, in the human IDR-ome (Figure 7b). The highest *f_C_shape_* scores occur for neutral chains (NCPR ∼ 0) with *ν* ∼ 0.5, indicating that high *f_C_shape_* scores are found for sequences and structures resembling GW chains (Figures 7b and 7c). As |NCPR| increases, *ν* tends to increase and *f_C_shape_* tends to decrease, likely because the increased charge densities restrict conformational diversity (compare to PEI at high protonation states, Figure 2). At NCPR ∼ 0, the IDRs exhibit a wide range of *ν* (ranging from ∼ 0.1 to ∼ 0.6 as well as *f_C_shape_* (from ∼ 0.5 to ∼ 0.85) (Figure 7b).

## Discussion

In this study, we introduce a polymer physics-based approach to examine the conformational landscapes of intrinsically disordered proteins and use this paradigm to study the human IDR-ome, a dataset previously published by Tesei *et al.*^19^ Specifically, we show that (RSA, *R_s_*) provides a universal framework for disordered chains, allowing us to resolve structural heterogeneity in IDRs, a hallmark of IDRs that is overlooked through the routine usage of ensemble-averaged metrics. Ensemble-averaged descriptors typically suffice for stably-folded proteins. However, IDRs can visit drastically different conformations, and using ensemble-averaged metrics may limit understanding of their sequence-structure-function relationships. To address this limitation, we use a joint distribution of two shape descriptors (RSA and *R_s_*), capturing conformational diversity through a two-dimensional representation. We apply this representation to the entire chain (global) and to subchains (local), revealing conformational heterogeneity both globally and locally. For instance, we show two IDRs (ZC3H4 and GCNA; Figures 5c and 5d, respectively) with similar *ν* but very different (RSA, *R_s_*) plots, which are traced to differences in their respective charge patterning (Figures 5g and 5h). We also show two IDRs (PYDC5 and BRPF3, Figure 6) that have similar global (RSA, *R*_s_) plots, but their subchain conformations differ significantly, which is likely driven by local sequence features.

Earlier, Tesei *et al.* showed^19^ that globally compact IDRs were correlated with specific protein functions and localization to cellular components. Additionally, chain compaction could indicate intramolecular attraction and may be associated with a propensity to phase separate, which could be biologically consequential.^19,48,49^ With subchain analysis, we identify globally non-compact IDRs with local compaction (subgroup 3), that are functionally similar to globally compact IDRs (subgroup 1) based on GO enrichment analysis. Further, subgroup 1 and subgroup 3 are associated with similarly high propensities for liquid-liquid phase separation (〈*p*_*LLPS*_〉). This strongly suggests that locally compact (but globally non-compact) IDRs are comparable in biological significance to globally compact IDRs.

A major objective of studies of IDR structures has been to link structure with both the underlying amino acid sequence and the cellular function of the protein. Ensemble-averaged structural descriptors of IDRs have previously been linked to sequence features, such as SHD, and function.^19^ However, the relationships between sequence features and the conformational heterogeneity of IDRs, particularly at the instantaneous level, remain largely unexplored. The (RSA, *R_s_*) scatter plots and heatmaps introduced here provide a framework to probe these relationships, as the two-dimensional representations of instantaneous conformations contained in these plots could potentially be predicted from sequence features using advanced deep learning methods. Since high conformational heterogeneity is a defining feature of IDRs, linking sequence, structure and function through our proposed framework will enhance understanding of their sequence-structure-function relationships.

Our framework for characterizing IDR conformations can also be potentially achieved experimentally, as end-to-end distance distributions (*R_ee_*) can be measured by fluorescence resonance energy transfer (FRET), while the hydrodynamic radius (*R_h_*, a quantity related to *R_g_*) can be determined using dynamic light scattering (DLS). As IDPs/IDRs are susceptible^50,51^ to regulation through protein phosphorylation, future experimental work could potentially pair polymer physics quantities to probe the phosphorylation-induced changes in conformational landscapes.

## Data Availability

This study re-analyzes a publicly-available dataset of human IDR simulations generated by Tesei *et al.*^19^ This study also re-uses IDR sequence and structural features computed and published by Tesei *et al.* A file (*Extended_Data_1.csv*), containing additional properties of those IDRs computed in this study, is provided.

## Supplemental Files

*Extended_Data_1.csv* – Properties of the IDRs in the human IDR-ome computed in this study *Supplementary_Data_1.xlsx* – Fractions of IDRs in subgroups 1, 2, 3 and 4 annotated with the GO terms (MF) that Tesei *et al.* found to be associated with more compact/extended IDRs

*Supplementary_Data_2.xlsx* – Fractions of IDRs in subgroups 1, 2, 3 and 4 annotated with the GO terms (CC) that Tesei *et al.* found to be associated with more compact/extended IDRs

*Supplementary_Data_3.xlsx* – Results of GO-term enrichment analyses (Molecular Function and Cellular Component) for proteins corresponding to IDRs in subgroups 1, 2, 3, and 4, relative to their respective background datasets (FDR-adjusted p < 0.05)

## Code Availability

*PyHeteroMap* (https://github.com/hshadman/IDP_Global_Local_Conformational_Landscapes) is a python package that can generate global (RSA, *R_s_*) scatter plots with *f_C_shape_* scores, subchain plots of 〈RSA〉, 〈*R*_*s*_〉, *ν* and other polymer properties, and (RSA, *R_s_*) scatter plots for one or more user-selected subchains, directly from IDR trajectory. Additionally, *PyHeteroMap* can simulate new Gaussian Walk (GW) chains (user provides the number of monomers of the chain and the number of snapshots of simulation).

## Acknowledgements

The authors wish to thank Professor Kresten Lindorff-Larsen (University of Copenhagen) for fruitful discussions.

## Author Contributions

J.D.Z., H.S., and Y.W. conceived ideas and designed the project. H.S. conducted all analysis, wrote code, and prepared all figures and tables, with input from Y.W., J.D.Z., Q.C., and M.L. H.S. conducted GW and SAW simulations with input from Y.W., J.D.Z., Q.C., and M.L. H.S. prepared the first draft of manuscript and H.S., J.D.Z., Q.C., M.L. and Y.W. revised the manuscript.

