## Supplementary Materials for "Resolving Local and Global Conformational Heterogeneity of the Human Intrinsically Disordered Proteome"

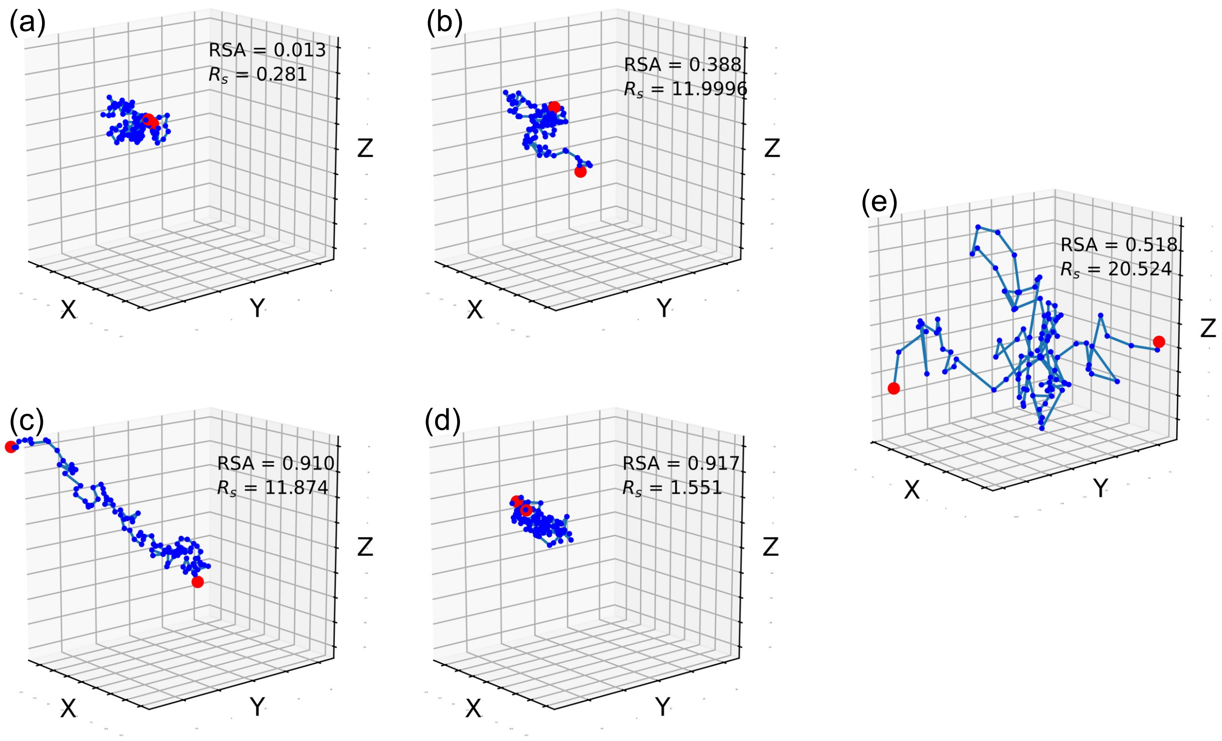


Figure S1: Images of a Gaussian Walk (GW) polymer chain conformation at relatively (a) low $R_s$, low RSA, (b) $R_s$ close to 12, low RSA, (c) $R_s$ close to 12, high RSA, (d) low $R_s$, high RSA, and (e) $R_s > 12$. Red represents terminal 'beads' (monomers), with all other monomers colored in blue.

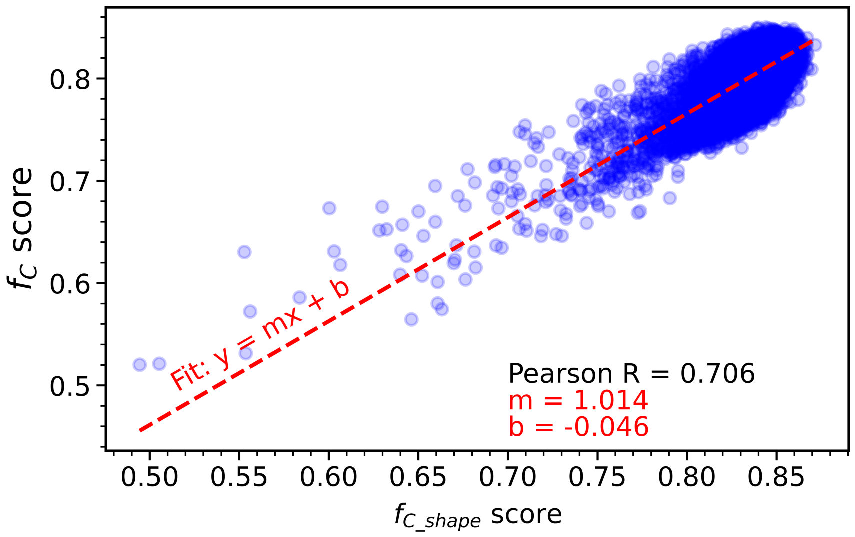


Figure S2: Scatter plot of $f_C$ against $f_{C\_shape}$ for the human IDR-ome dataset

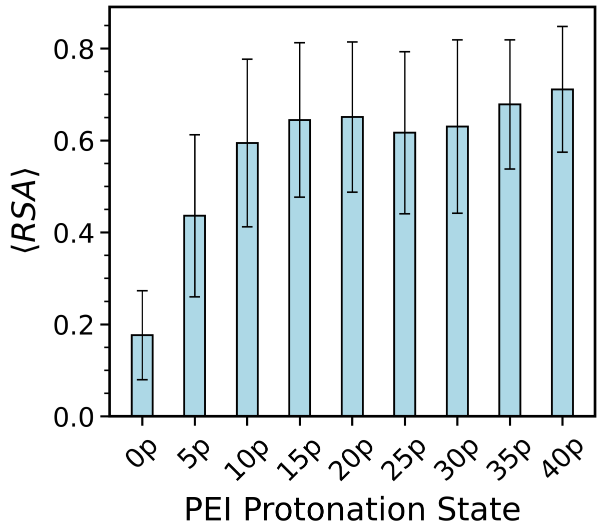


Figure S3: Mean RSA at each protonation state of PEI, with error bars showing standard deviation

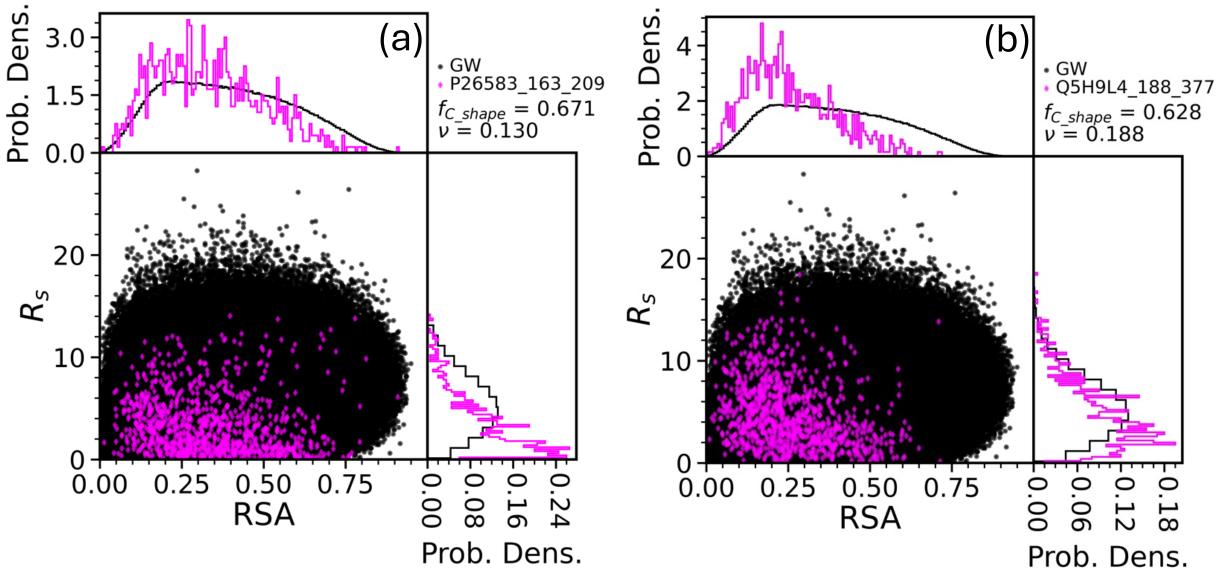


Figure S4: Scatter plots of $R_s$ against RSA for two IDR simulations, against a GW reference. (a) and (b) have low ν.

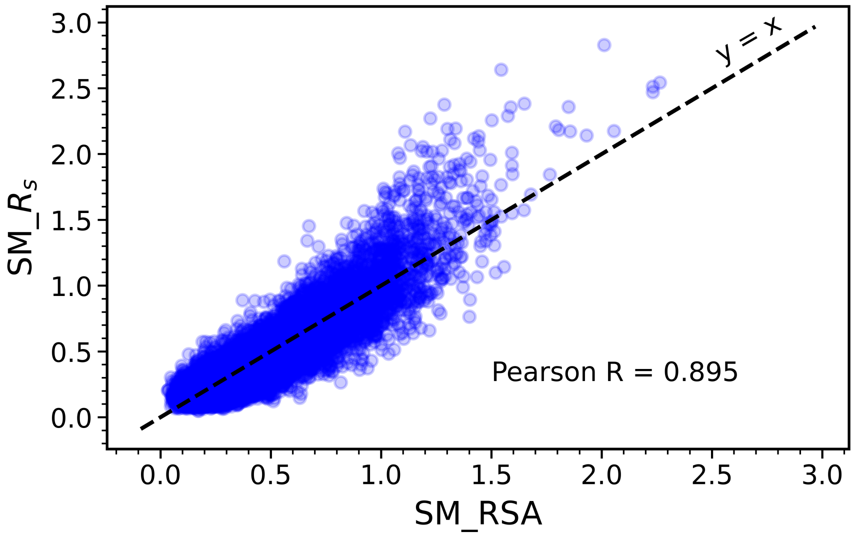


Figure S5: A scatter plot of SM_$R_s$ against SM_RSA for all ~28000 IDRs in the IDR-ome dataset.

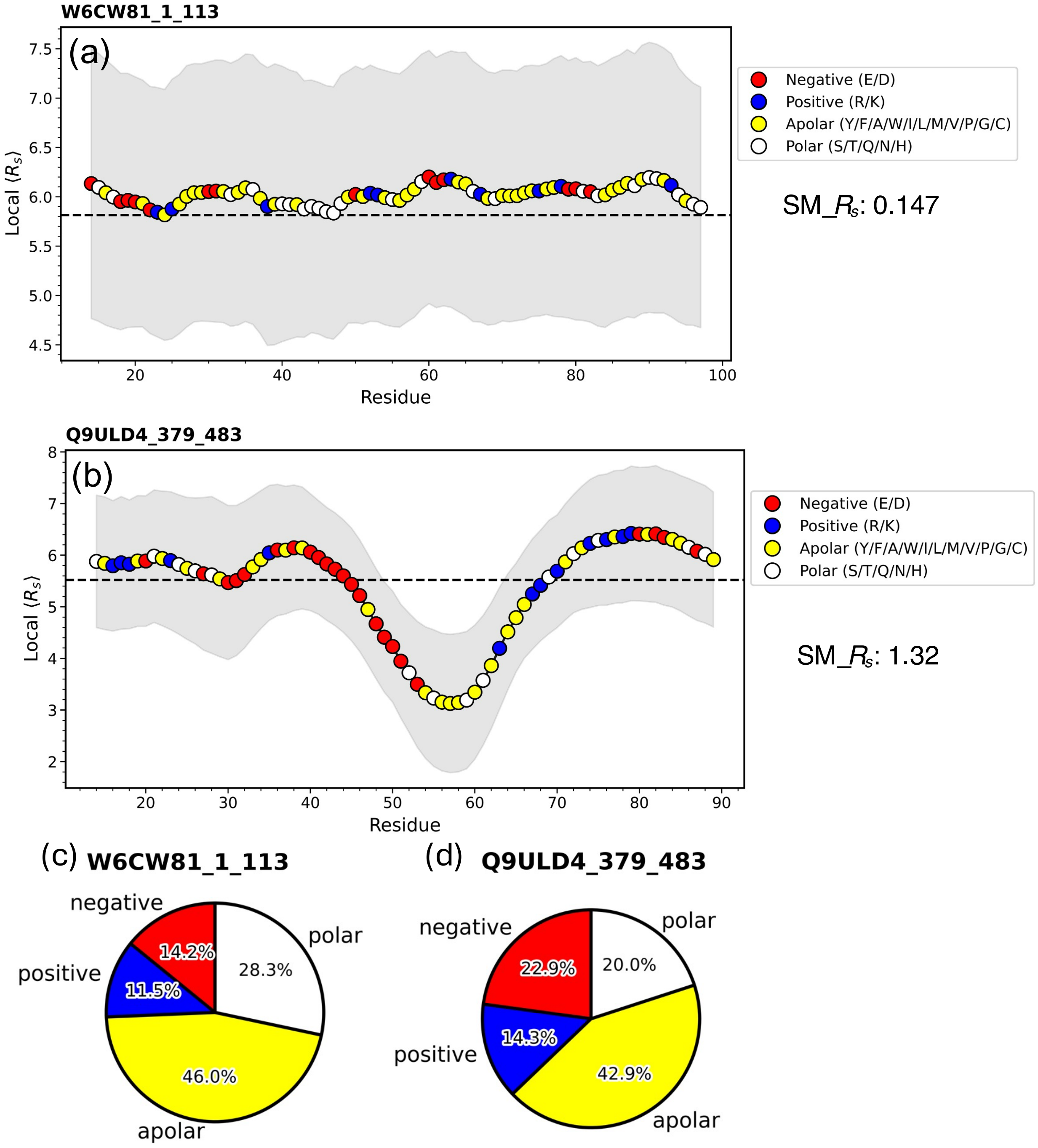


Figure S6: (a) and (b): Local variation of $\langle R_s \rangle$ for IDR chains (identity shown in bold). The gene name for the protein corresponding to the IDR in (a) is PYDC5 and (b) is BRPF3. For (a) and (b), the x-axis identifies middle residue for each moving window, the grey region shows the standard deviation, and the dashed line identifies $\langle R_s \rangle$ for the whole chain (global). (c) and (d) show amino acid sequence compositions for the IDRs from PYDC5 and BRPF3, respectively.

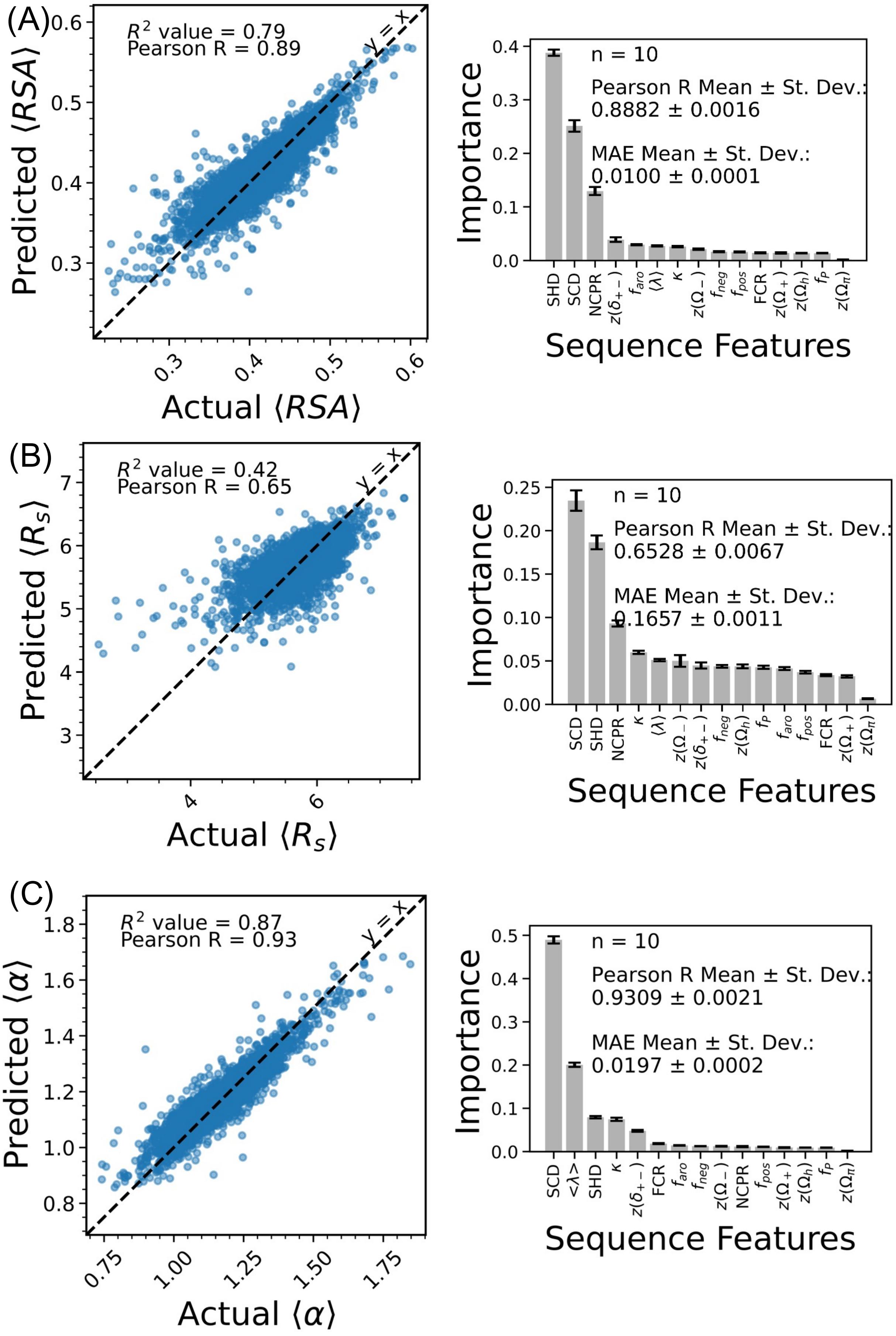
(A)
$R^2$ value = 0.79
Pearson R = 0.89
y = x
Predicted ⟨RSA⟩
Actual ⟨RSA⟩
n = 10
Pearson R Mean ± St. Dev.:
0.8882 ± 0.0016
MAE Mean ± St. Dev.:
0.0100 ± 0.0001
Importance
Sequence Features
SHD SCD NCPR $z(\delta_{+-})$ $f_{aro}$ $\langle\lambda\rangle$ $\kappa$ $z(\Omega_-)$ $f_{neg}$ $f_{pos}$ FCR $z(\Omega_+)$ $z(\Omega_h)$ $f_P$ $z(\Omega_\pi)$
(B)
$R^2$ value = 0.42
Pearson R = 0.65
y = x
Predicted $\langle R_s \rangle$
Actual $\langle R_s \rangle$
n = 10
Pearson R Mean ± St. Dev.:
0.6528 ± 0.0067
MAE Mean ± St. Dev.:
0.1657 ± 0.0011
Importance
Sequence Features
SCD SHD NCPR $\kappa$ $\langle\lambda\rangle$ $z(\Omega_-)$ $z(\delta_{+-})$ $f_{neg}$ $z(\Omega_h)$ $f_P$ $f_{aro}$ $f_{pos}$ FCR $z(\Omega_+)$ $z(\Omega_\pi)$
(C)
$R^2$ value = 0.87
Pearson R = 0.93
y = x
Predicted $\langle\alpha\rangle$
Actual $\langle\alpha\rangle$
n = 10
Pearson R Mean ± St. Dev.:
0.9309 ± 0.0021
MAE Mean ± St. Dev.:
0.0197 ± 0.0002
Importance
Sequence Features
SCD <λ> SHD $\kappa$ $z(\delta_{+-})$ FCR $f_{aro}$ $f_{neg}$ $z(\Omega_-)$ NCPR $f_{pos}$ $z(\Omega_+)$ $z(\Omega_h)$ $f_P$ $z(\Omega_\pi)$

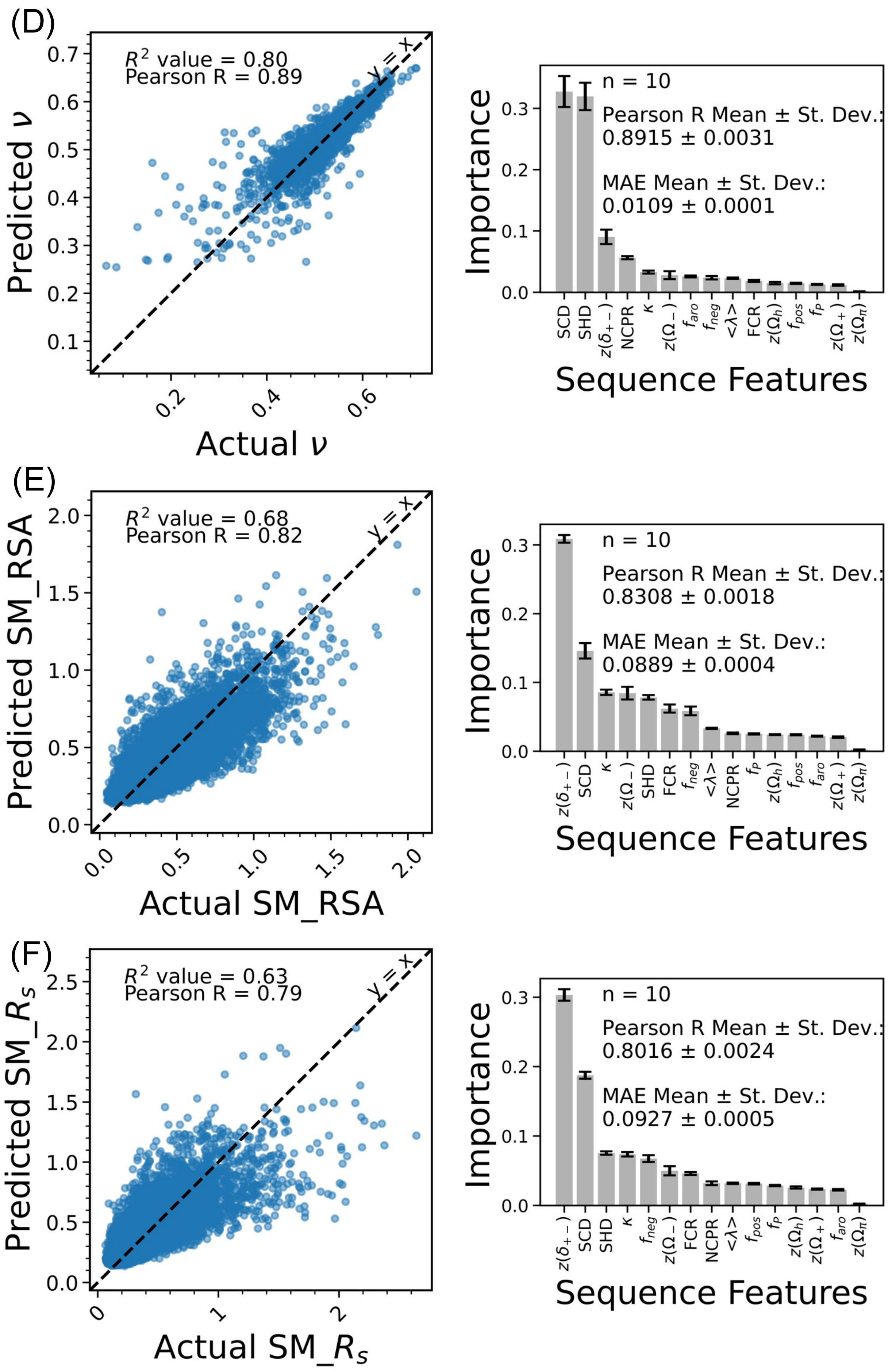


Figure S7: Predictions of different descriptors using random forest regression models based on sequence features for all ~28000 human IDRs. For each panel, the left figure shows actual vs predicted values for a single test set, and the right figure shows the feature importance of the sequence features (mean ± standard deviation of 10 different random train-test splits).

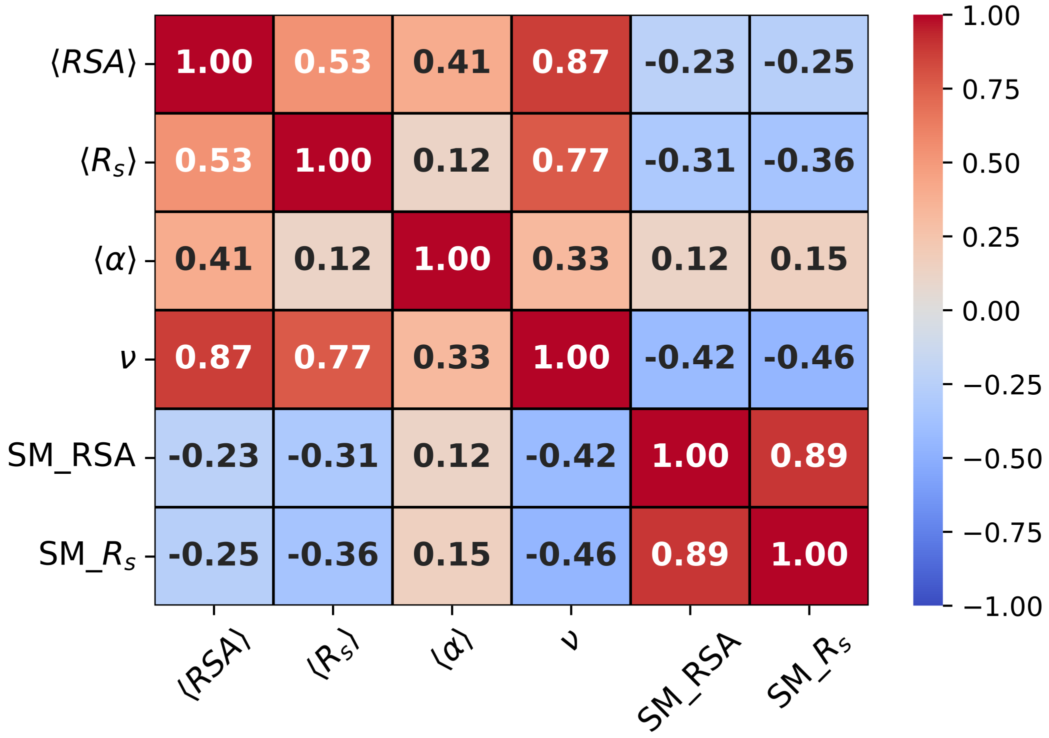


Figure S8: A correlation matrix of Pearson R values between different conformational properties and SM quantities for the human IDR-ome dataset

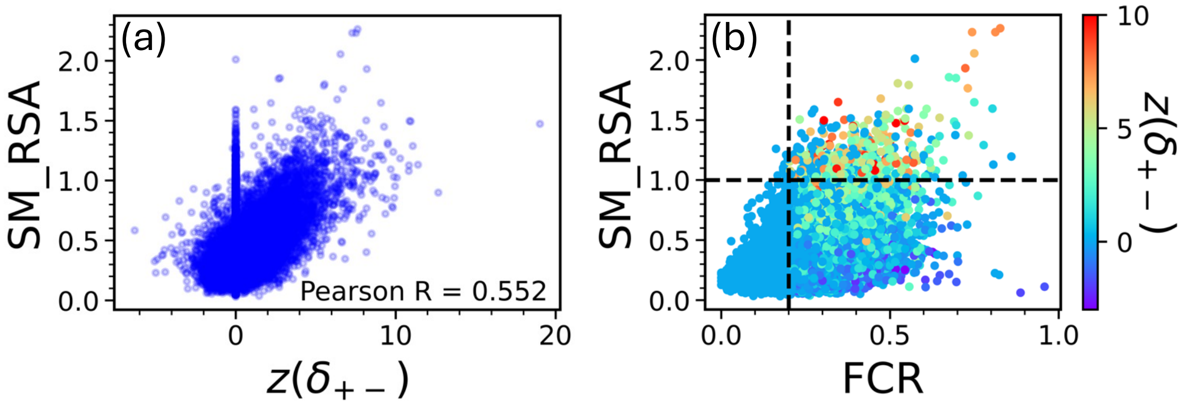


Figure S9: Scatter plots of (a) SM_RSA against $z(\delta_{+-})$ and (b) SM_RSA against fraction of charged residues (FCR) colored by $z(\delta_{+-})$, for all ~28000 IDRs in the human IDR-ome dataset.

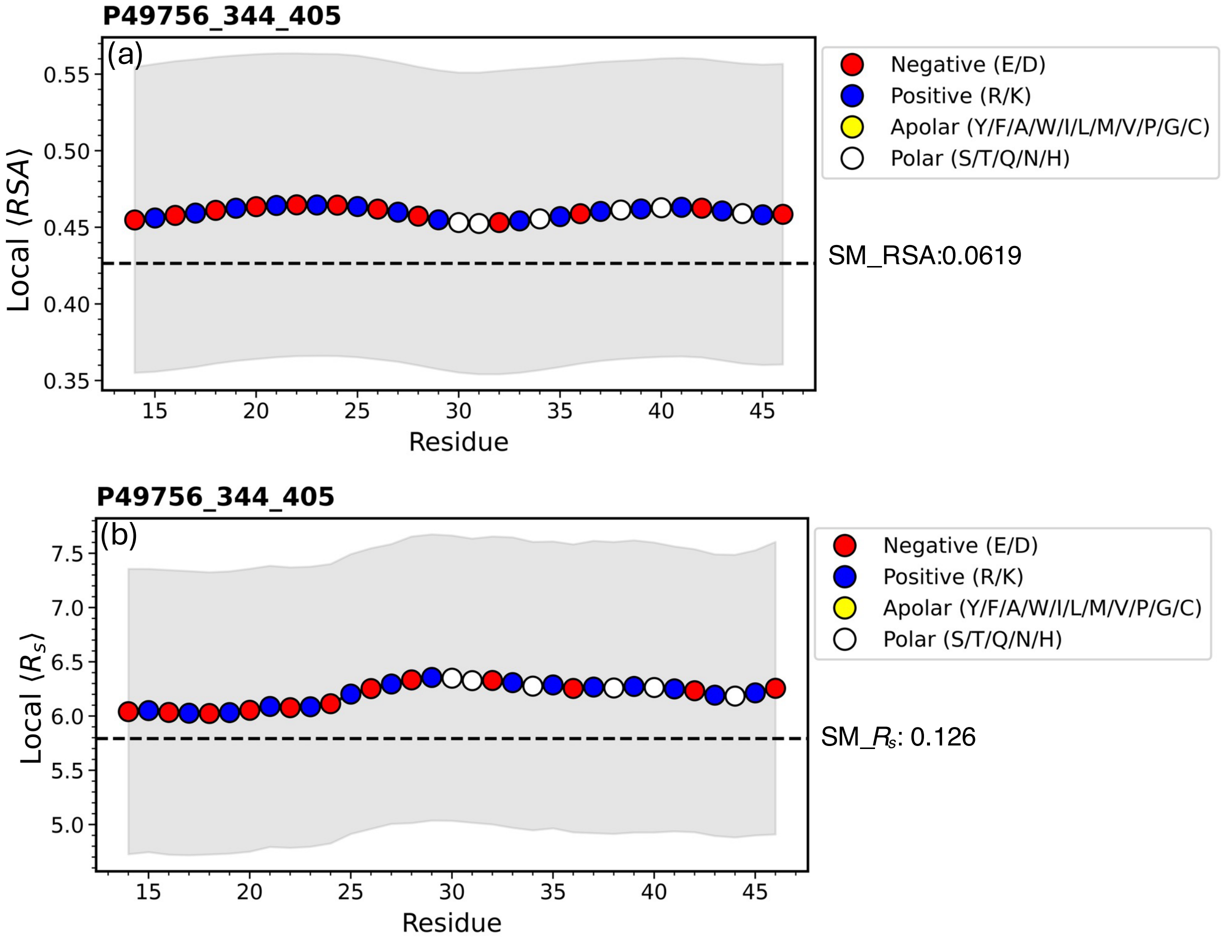


Figure S10: Local variation of (a) $\langle RSA \rangle$ and (b) $\langle R_s \rangle$, across an IDR (identity shown in bold) chain. X-axis identifies middle residue for each moving window, the gray region shows standard deviation, and the dashed line identifies the whole chain (global) value. This sequence has FCR 0.887, and $z(\delta_{+-})$ -1.827. The gene name for the protein associated with this IDR is RBM25.

Table S1: Subgroups of IDRs used for GO enrichment analysis

| | Condition* | Number of IDRs of Subgroup in MF List⊗ | Number of Unique Proteins in MF List+ | Number of IDRs of Subgroup in CC List⊗ | Number of Unique Proteins in CC List+ |
|---|---|---|---|---|---|
| Subgroup 1 | $\nu < 0.45$ | 479 | 465 | 550 | 533 |
| Subgroup 2 | $\nu > 0.55$ | 7322 | 5559 | 8270 | 6214 |
| Subgroup 3 | $\nu \geq 0.45$ and $\nu_{min\_local} < 0.45$ (within the same IDR) | 1163 | 1096 | 1261 | 1187 |
| Subgroup 4 | $\nu \leq 0.55$ and $\nu_{max\_local} > 0.55$ (within the same IDR) | 15092 | 9656 | 17041 | 11004 |

*$\nu$ is the whole chain (global) Flory scaling exponent of the IDR; $\nu_{min\_local}$ is the minimum local (subchain) $\nu$ for the IDR, $\nu_{max\_local}$ is the maximum local (subchain) $\nu$ for the IDR.

+This counts the number of unique proteins associated with the IDRs in each MF or CC subgroup. Subgroups 1 and 3 in the MF category share 54 unique proteins. Subgroups 2 and 4 in the MF category share 3312 unique proteins. Subgroups 1 and 3 in the CC category share 59 unique proteins. Subgroups 2 and 4 in the CC category share 3678 unique proteins.

⊗The MF list has 22,445 IDRs in total and the CC list has 25,364 IDRs in total.

Table S2: Subgroups of IDRs and their $\langle p_{LLPS} \rangle$ values

| | Number of IDRs in subgroup∀ | Number of unique proteins in subgroup | Number of unique proteins in subgroup with available $p_{LLPS}$ values | $p_{LLPS}$ (mean ± standard deviation) |
|---|---|---|---|---|
| Subgroup 1 | 597 | 580 | 580 | $0.76 \pm 0.28$ |
| Subgroup 2 | 8896 | 6743 | 6740 | $0.54 \pm 0.32$ |
| Subgroup 3 | 1439 | 1362 | 1353 | $0.76 \pm 0.28$ |
| Subgroup 4 | 19102 | 12626 | 12606 | $0.62 \pm 0.32$ |

∀$p_{LLPS}$ values correspond to the unique proteins containing the IDRs, not the IDRs alone. The IDR subgroups defined here are from the main human IDR-ome dataset containing 28,054 IDRs (see *Methods*), using the same conditions defined in Table S1. Subgroups 1 and 3 share 63 unique proteins. Subgroups 2 and 4 share 3980 unique proteins. The $p_{LLPS}$ values corresponding to the UniProt IDs of the proteins in the human proteome have been published previously (see *Methods*).

**Extended_Data_1.csv (separate file)**– Properties of the IDRs in the human IDR-ome computed in this study

**Supplementary_Data_1.xlsx (separate file)** – Fractions of IDRs in subgroups 1, 2, 3 and 4 annotated with the GO terms (MF) that Tesei et al. found to be associated with more compact/extended IDRs

**Supplementary_Data_2.xlsx (separate file)** – Fractions of IDRs in subgroups 1, 2, 3 and 4 annotated with the GO terms (CC) that Tesei et al. found to be associated with more compact/extended IDRs

**Supplementary_Data_3.xlsx (separate file)** – Results of GO-term enrichment analyses (Molecular Function and Cellular Component) for proteins corresponding to IDRs in subgroups 1, 2, 3, and 4, relative to their respective background datasets (FDR-adjusted $p < 0.05$)
